# Long-Read Transcriptome Sequencing and Functional Validation Reveals Novel and Oncogenic Gene Fusions in Fusion Panel-Negative Gliomas

**DOI:** 10.64898/2026.03.13.711117

**Authors:** Karleena Rybacki, Emily Na Young Cha, Hannah M. Deutsch, Eloise Gaudet, Mian Umair Ahsan, Feng Xu, Joe Chan, Marilyn Li, Yuanquan Song, Kai Wang

## Abstract

Gliomas comprise a heterogeneous group of central nervous system tumors in which gene fusions (GFs) are significant oncogenic drivers and emerging diagnostic and therapeutic biomarkers. In cancer diagnosis, GF detection largely relies on targeted short-read sequencing fusion panels, such as the Children’s Hospital of Philadelphia (CHOP) Fusion Panel (FUSIP). While these panels are effective for detecting recurrent, well-characterized GFs, they are limited to predefined gene sets and cannot identify full-length transcripts. Here, we analyzed 49 high-and low-grade gliomas previously classified as fusion-negative by FUSIP using an untargeted whole-transcriptome RNA sequencing approach with Oxford Nanopore Technologies (ONT) long-read sequencing. This enabled transcriptome-wide fusion discovery of additional known and potentially novel oncogenic GFs beyond panel constraints. Long-read sequencing further allowed direct resolution of full-length fusion transcripts and their associated isoform structures. By integrating GF detection with isoform-level transcript analysis, we identified fusion-associated transcript isoforms with alternative splicing patterns that aligned near reported GF breakpoints, including *ZNF254*::*GNAS* and *PTPRK*::*NOX3*, which have not been reported in literature or existing fusion databases. To assess functional relevance, candidate GFs were evaluated using the *Drosophila melanogaster* model, with ventral nerve cord (VNC) morphology serving as a quantitative *in vivo* readout of fusion-induced disruption of glial regulation. VNC enlargement or elongation reflects abnormal glial growth or defects in brain tissue organization. Of the 15 candidate GFs subjected to experimental functional testing, 8 induced significant VNC abnormalities relative to wild-type controls, indicating fusion-specific disruption and oncogenic potential. Notably, *CLDND1*::*WRN* and *DUSP22*::*APOE* produced the most pronounced VNC phenotypes. Together, these findings demonstrate that untargeted transcriptome-wide GF discovery, coupled with long-read isoform-level analysis and *in vivo* functional validation, enables the identification and prioritization of potentially novel and clinically relevant GFs that are missed by standard targeted short-read fusion panels in glioma.

## INTRODUCTION

Gliomas are a diverse group of brain tumors that originate from glial cells of the central nervous system (CNS), accounting for approximately 23% of all primary brain tumors and 81% of malignant brain tumors.^1^ These tumors arise in different glial cell types and vary in location, histology, growth pattern, and genetic profile, which are critical for understanding their clinical prognosis.^1,2^ Collectively, these features provide valuable insights that can be utilized to stratify patient risk, predict prognosis, and guide treatment selection. The World Health Organization (WHO) classifies gliomas from grades I to IV primarily based on histological and molecular features. Grades I and II attribute to low-grade gliomas (LGGs), grades III and IV for high-grade gliomas (HGGs), and glioblastoma multiforme (GBM) being one of the most aggressive grade IV subtypes.^3^ LGGs are slow-growing with a more favorable prognosis, whereas HGGs are aggressive cancers and present therapeutic challenges due to resistance to conventional therapies.^4,5^ In addition, standard treatments for gliomas, including surgical resection, radiotherapy, and chemotherapy, are often associated with substantial adverse effects and limited efficacy in aggressive disease scenarios, highlighting the need for more precise and less toxic therapeutic strategies.^6,7^ This diverse clinical range, driven by various genomic alterations, including gene fusions (GFs), underscores the need for comprehensive analysis approaches to enhance precision diagnostics.^8,9^

The advent of next-generation sequencing has significantly advanced the diagnosis and classification of glioma by incorporating recurrent genetic alterations through targeted enrichment methods. Foundational glioma biomarkers routinely used in diagnostics include isocitrate dehydrogenase (IDH) mutations, 1p/19q co-deletion, *EGFR* gene amplification, and the gain of chromosome 7 with the concurrent loss of chromosome 10.^10–13^ Additionally, structural variants including GFs and transcript isoform alternative splicing events have been identified as important contributors to the molecular complexity of glioma.^14–16^ These genomic rearrangements can lead to aberrant gene expression, chimeric transcripts, or disruption of tumor suppressor pathways. In glioma, recurrent GFs like *FGFR3::TACC3*, *KIAA1549::BRAF*, and *EGFR::SEPT14*, and fusions involving *ALK*, *NTRK*, *ROS1*, and *MET*, have been recognized as promising targets for personalized treatment options.^16–19^ Notably, *NTRK* fusions have emerged as promising therapeutic targets in solid tumors, including glioma, with FDA-approved therapies such as entrectinib and larotrectinib.^20^ Ongoing clinical trials are further evaluating the use of larotrectinib in pediatric patients with *NTRK* fusion-positive HGGs.^21^

Clinical methods for detecting chromosomal alterations still rely heavily on targeted approaches that are limited to a predefined list of genes. This inherently restricts the ability to identify potentially novel or complex alterations that may have diagnostic or therapeutic relevance. For example, short-read sequencing fusion panels, such as the Children’s Hospital of Philadelphia (CHOP) Fusion Panel (FUSIP), are designed to detect known GFs within a curated set of 119 oncogenes implicated in cancer-related GFs.^22^ While these panels are efficient and effective for detecting recurrent, clinically actionable GFs, they are constrained by several technical limitations. Short reads (typically 150-300 bps) cannot span full-length transcripts, making it difficult to resolve GF structures or complex rearrangements. Additionally, alignment challenges in repetitive or highly complex regions of the genome may result in ambiguous or missed GF calls. Most critically, limiting the detection to a predefined set of genes risks overlooking potentially novel or rare fusions that may be clinically relevant. These limitations are particularly evident in diagnostically challenging cases where standard short-read approaches fail to identify oncogenic rearrangements, despite clinical evidence suggestive of their presence, potentially leaving actionable GFs undetected. These technical challenges highlight the need for more comprehensive sequencing and analyses that can overcome the inherent limitations of targeted, short-read approaches.

Long-read sequencing technologies, such as Pacific Biosciences and Oxford Nanopore Technologies (ONT), facilitate the rapid generation of longer reads (typically >10kb) that span entire transcripts. Recent improvements in ONT technology have reduced error rates to approximately 1%, further enhancing its accuracy.^23^ By capturing full-length transcripts, ONT enables precise characterization of fusion breakpoints without the need for computational read reconstruction inherent to short-read approaches. This allows for the discovery of potentially novel GFs and splice variants that may contribute to oncogenesis. In our previous study, we established a long-read sequencing-based GF detection and analysis pipeline that integrated multiple long-read fusion callers with downstream computational filtering to annotate and prioritize candidate GFs.^24^ In this glioma cohort, we found that long-read sequencing could recover GFs involving a gene targeted in FUSIP and also revealed additional candidate GFs not captured by the panel or represented in fusion databases such as COSMIC Fusion^25^, Mitelman DB^26^, and ChimerDB 4.0^27^. However, this unbiased whole-transcriptome approach also generated many putative GFs whose biological and clinical relevance was unknown. Without systematic prioritization and functional validation, it is challenging to distinguish true oncogenic fusions from rare passenger or artifactual events. Building upon this framework, we expanded our original cohort to a total of 49 glioma samples that were previously classified as fusion-negative by the CHOP FUSIP. These negative cases are diagnostically challenging, as disease drivers could have been missed by targeted short-read methods. Long-read sequencing is particularly well suited for such cases, as it directly resolves full-length transcripts and enables unbiased detection of GFs and complex rearrangements that targeted approaches may overlook.

To functionally assess the oncogenic potential of these candidate fusions, we incorporated *in vivo* functional assays using *Drosophila melanogaster*, fruit flies, to validate and contextualize our computational GF predictions. *Drosophila* share many conserved genes and signaling pathways with humans, and has been widely used as a robust tumor model for identifying genetic drivers of brain tumorigenesis and other neurological diseases.^28–32^ In general, the fruit fly is a valuable model organism to functionally analyze GFs due to their short lifecycle and genetic accessibility.^29^ By leveraging the GAL4/UAS system to induce glia-specific overexpression of individual GFs, we evaluated their effects on cell proliferation and neural development within the CNS. In this model, the ventral nerve cord (VNC), which is a structure analogous to the human spinal cord, serves as a quantitative readout of neural/glial overgrowth, where VNC enlargement or elongation suggests oncogenicity. Functional genomic assays such as these offer deeper insights into the biological consequences of rare or potentially novel GFs not identified by existing diagnostic approaches. Prior studies have demonstrated that systematic functional characterization of GFs can uncover key oncogenic drivers with therapeutic potential.^33,34^ By integrating whole-transcriptome long-read GF detection, computational prioritization, and experimental functional validation, our approach aims to identify rare or novel oncogenic GFs in gliomas to uncover underlying biological mechanisms towards improving precision diagnostics in glioma.

## RESULTS

We applied ONT long-read sequencing to 49 FUSIP panel-negative gliomas (24 LGG and 25 HGG) to comprehensively characterize GFs and transcript isoforms across a diverse set of gliomas, including glioblastomas, IDH-mutant astrocytomas, oligodendrogliomas, diffuse midline gliomas, and pilocytic astrocytomas, with all clinical indications summarized in **Supplemental Table S1**. As outlined in **Figure 1**, this approach enabled unbiased GF detection, experimental validation, and functional prioritization of candidate GFs. In parallel, long-read sequencing also provided full-length transcript resolution, allowing us to identify alternative isoform expression patterns in the fusion panel-negative cohort. Prioritized GFs expressed in the *Drosophila* glial cells produced GF-driven phenotypes, providing *in vivo* evidence of their oncogenic potential to identify candidate molecular drivers and potential therapeutic targets.

**Figure 1.**
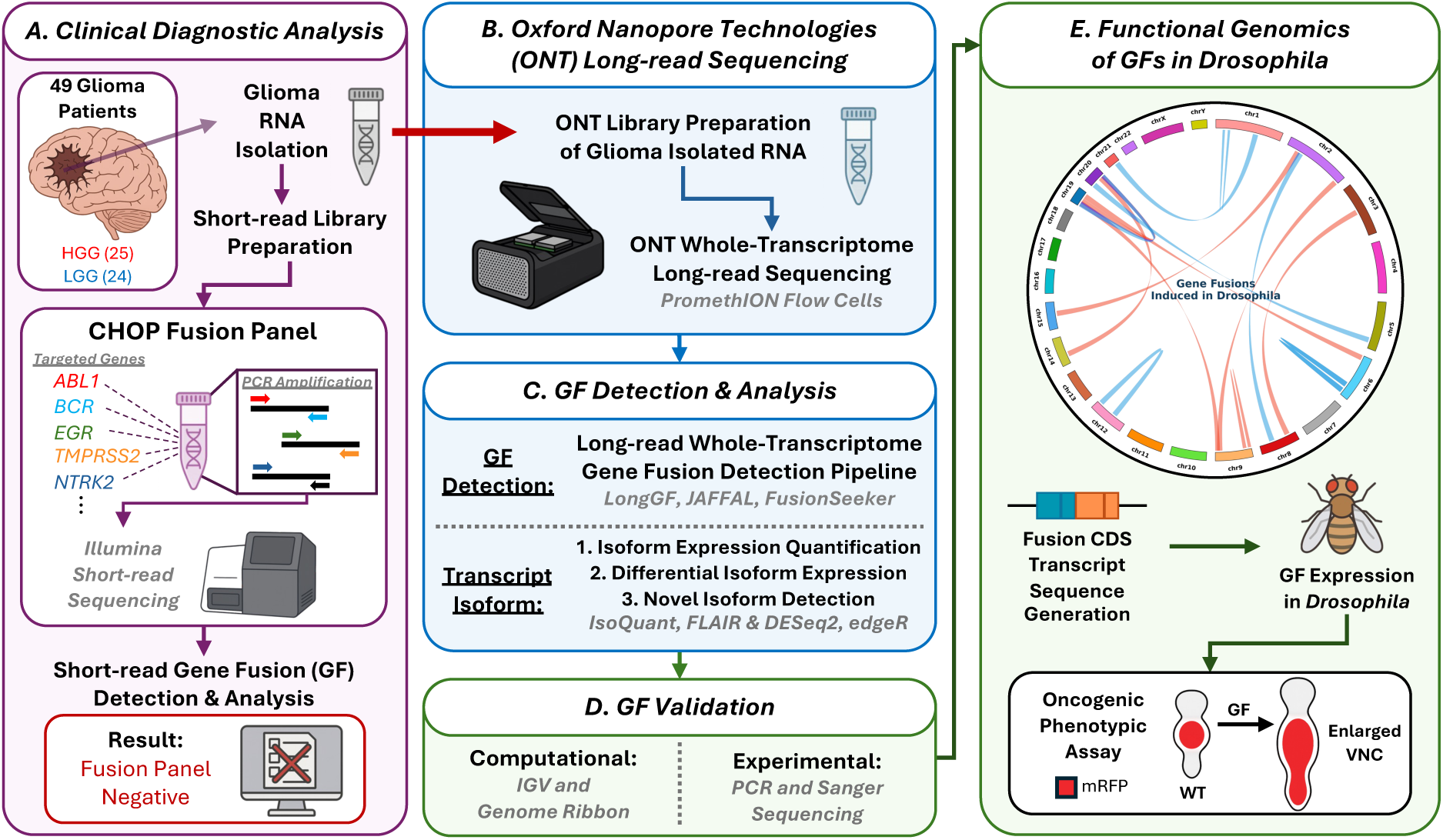
Integrated workflow of GF identification, validation, and functional assessment in fusion panel-negative glioma samples. **(A)** Clinical diagnostic testing was previously performed on the 49 glioma patients (25 HGG, 24 LGG) using the short-read CHOP Fusion Panel. The panel testing was negative for known clinically relevant GFs. **(B)** To enable more comprehensive GF discovery, ONT long-read whole-transcriptome sequencing was performed on RNA isolated from the same glioma samples. **(C)** Downstream analysis included GF detection using the long-read whole-transcriptome GF detection pipeline and orthogonal transcript isoform analyses. **(D)** Prioritized candidate GFs were subjected to both computational and experimental validation. **(E)** Successfully validated GFs (displayed on the Circos plot, with connections of GFs from HGG samples in red, and LGG samples in blue) were functionally assessed by in vivo fusion transcript expression in *Drosophila melanogaster* tumor models. Resulting oncogenic phenotypes (i.e. enlarged/shrunken VNC length or VNC area) were quantified to evaluate the biological impact of potentially novel, glioma-associated GFs.

### Long-read Sequencing Quality Metrics and Sample Characteristics

ONT PromethION long-read sequencing was performed across six sequencing pools, including 24 samples in our previous study^24^ and 25 newly sequenced samples. Each pool contains a mixture of LGG and HGG samples to reduce batch effects. For the newly sequenced glioma samples, read N50 values ranged from 265 to 1,164 bp, with an average of 3.43 Gb and 5.91 M reads per sample, and all libraries passed internal quality checks (**Supplemental Table S2**). Flow cell performance was monitored for up to 96 hours and confirmed that sufficient data for GF detection was obtained within 48-72 hours (**Supplemental Table S3**). After alignment to GRCh38, LongReadSum^35^ metrics across all 49 samples showed a mean of 6.2 M total reads per sample (range 1.6-21.2 M) and an average read length of 544 bp (range 186-1,001 bp), with most reads successfully mapped to the reference genome (**Supplemental Table S4**). Sequencing yield and metrics differed significantly between pools (one-way ANOVA, all p-values <0.05), but both HGG and LGG samples were distributed across pools and Mann-Whitney U tests confirmed no significant differences in sequencing metrics between glioma grades. All samples generated sufficient data to constitute a high-quality long-read dataset for glioma, enabling subsequent analyses for GF detection, transcript isoform analysis, and downstream functional studies.

### Gene Fusion Detection and Characterization

All 49 long-read sequenced glioma samples were processed through our previously established long-read whole-transcriptome GF detection pipeline, with additional downstream filters applied as described in the Methods. This approach integrates multiple GF detection tools such as LongGF, JAFFAL, and FusionSeeker. Initial ensemble fusion calling identified 110,969 unique GFs across the cohort prior to filtering. Here, unique GFs refer to distinct gene pairings, where a fusion and its reciprocal within 50 bp were treated as the same event within a sample (see Methods). This large initial number reflects the expected combined output of unbiased whole-transcriptome GF detection prior to any biological or clinical prioritization. Through a multistep filtering strategy to remove recurrent technical artifacts, non-expressed or non-coding events, fusions observed in healthy brain tissue, and fusions lacking cancer-relevant gene partners, we reduced this set to 1,249 unique high-confidence GFs across the cohort. This reduction reflects deliberate prioritization rather than permissive retention, yielding an average of 28 unique candidate GFs per sample (**Supplemental Table S5**), comparable to typical whole-transcriptome GF studies.

When restricting the analysis to fusions involving genes targeted by the CHOP FUSIP, only 175 unique candidate GFs were identified across the cohort. Expanding the analysis to include genes from the COSMIC Cancer Gene Census increased the number of unique candidate GFs by 6.8-fold (from 175 to 1,186), demonstrating that whole-transcriptome profiling beyond a targeted gene panel may substantially increase GF discovery in this cohort. Most samples (41/49, 83.7%) had fewer than 50 fusions remaining after filtering, whereas two samples (#34 and #40) had more than 200 fusions, and four samples (#71, #76, #78, and #79) had no fusions remaining, potentially reflecting both biological heterogeneity and technical variation in sequencing depth and quality. Across all unique fusions, the median support was 3 reads (mean 4.8 reads per fusion). A subset of fusions had been previously reported in public fusion databases, suggesting potential clinical relevance; for example, we identified the reciprocal of the *SOD1*::*TYK2* fusion (sample #63), detected by two GF tools, which has been previously reported in the Mitelman Database but not described in glioma.

To evaluate the functional potential of the remaining GFs after filtration, we annotated each breakpoint region using a three-tiered strategy that prioritized MANE Select transcripts, followed by the longest protein-coding transcript per gene, and finally all annotated transcripts (see Methods). Breakpoints within coding sequences (CDS) or intronic regions within 200 bp of exon boundaries (“near exon”) were classified as potentially functional. On average, 15.9% of GFs per sample (mean 6.8) met this CDS/near-exon criterion. These CDS/near exon breakpoints may represent alternative splice site usage, regions of microhomology that are challenging for GF detection tools to resolve precisely, or limitations of current transcript annotation databases. Fusion breakpoint classification revealed distinct patterns between the 5’ and 3’ genes in the GFs, where 42.4% of 5’ gene breakpoints fell in CDS/near exon regions compared to 30.6% for 3’ gene breakpoints, whereas 3’ genes had more 3’ UTR breakpoints (50.9% compared to 32.4% in the 5’ gene). Intronic and intergenic breakpoints were relatively rare for both gene partners (12.7% of all breakpoints). This annotation revealed that 97.1% of fusion breakpoints were successfully annotated using MANE Select transcripts, <1% required the longest protein-coding transcript, and 2.1% needed comprehensive searching across all transcripts. The GFs were largely grade-specific, with 804 fusions unique to HGG (67.8%), 362 unique to LGG (30.5%), and only 20 shared across grades (1.7%), although breakpoint distributions were broadly similar between HGG- and LGG-unique GFs.

We next assessed the biological and potential clinical relevance of GFs with potentially functional breakpoints (those that met the CDS/near-exon criterion) by annotating each GF against curated cancer and glioma gene lists. Of these potentially functional GFs, only 15.2% included at least one gene from CHOP FUSIP, indicating that several clinically recognized fusion partner genes were present despite the samples tested negative initially. Notably, one LGG fusion, *FUS*::*EGFR*, involved two panel genes that was not reported in any online fusion database. Among the functional GFs, 97% included a gene from the COSMIC Cancer Gene Census, and 8.3% involved a gene from a glioma-specific or CNS-associated gene list, suggesting that a subset of GFs involved genes previously implicated in glioma biology. Overall, 60.7% of fusions involved a known oncogene and 44.5% involved a tumor suppressor gene, with 3.8% (10 GFs) linking an oncogene to a tumor suppressor. A substantial fraction of GFs contained genes with high essentiality in glioma-related cell lines based on DepMap (46.2% with dependency probability >0.9 and 35.7% with gene effect score <-1), and 52.6% included genes previously reported as recurrent fusion partners in CNS tumors in the Mitelman Database. Collectively, these findings indicate that the prioritized high-confidence fusions in FUSIP-negative gliomas are enriched in cancer- and glioma-relevant genes, suggesting their biological and potential clinical relevance.

### Transcript Isoform Detection, Differential Expression, and Novel Isoform Characterization

To characterize transcript-level changes associated with glioma, we used IsoQuant and FLAIR to identify and quantify long-read transcript isoforms and performed differential expression (DE) analyses across four group comparisons: glioma vs. controls, HGG vs. controls, LGG vs. controls, and HGG vs. LGG (see Methods). Long-read whole transcriptome analysis of the glioma samples detected 47,023 unique transcript isoforms from IsoQuant and 23,670 from FLAIR, with 13,850 isoforms (24.4%) shared between the two tools. After expression-based filtering (see Methods), 6,902 IsoQuant and 2,797 FLAIR transcripts (2,157 shared) were retained for differential expression analysis. High-confidence DE isoforms were defined as those significant in both edgeR and DESeq2 (FDR or adjusted p-value ≤0.05 and the absolute value of the log2 FC in expression ≥1). This yielded the largest number of DE isoforms from the glioma vs. controls comparison (1,854 IsoQuant, 811 FLAIR) and the fewest in the HGG vs. LGG comparison (11 IsoQuant, 24 FLAIR), consistent with substantial transcriptomic differences between normal brain and glioma tissues but greater similarity amongst different grades of gliomas. Notably, only 19.4% of significant DE isoforms overlapped between IsoQuant and FLAIR, indicating that the two tools capture complementary aspects of the transcriptome and that their combined use expands overall isoform discovery. To summarize isoform-level patterns, consensus volcano plots are shown in **Figure 2A-B**, and **Supplementary Figure S2** shows concordance plots comparing edgeR and DESeq2 that highlight consistent patterns of differential transcript expression across all glioma comparisons.

**Figure 2.**
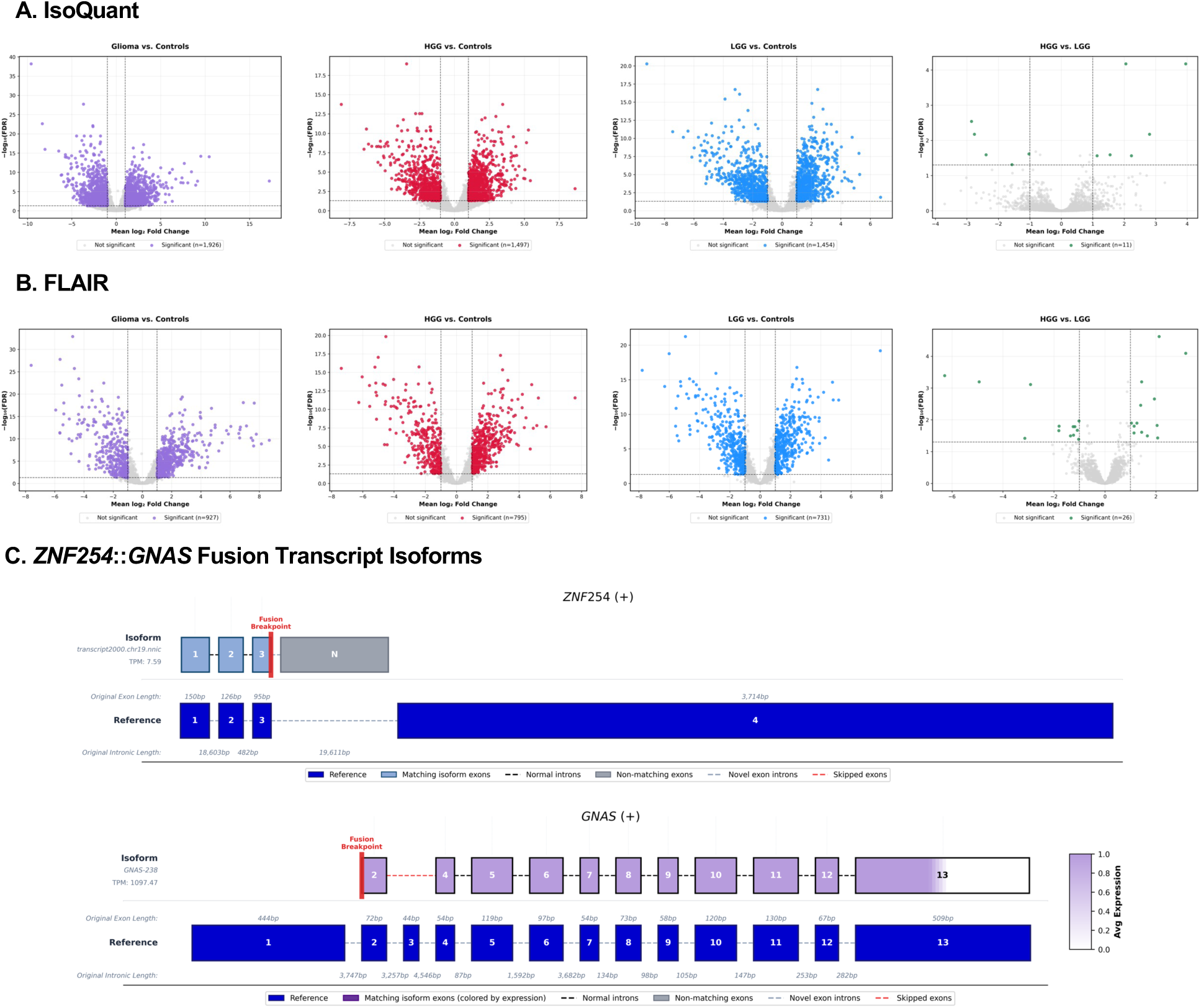
Differential transcript isoform expression across glioma comparisons and fusion transcript structures. **(A)** IsoQuant and **(B)** FLAIR consensus isoform volcano plots showing significantly differentially expressed transcript isoforms identified by both edgeR and DESeq2 across the remaining glioma sample comparisons with significantly DE isoforms: glioma vs. control, HGG vs. control, LGG vs. control, and HGG vs. LGG. Each point represents an individual isoform, with colored isoforms are significant and gray denotes non-significant isoforms. Gray dotted lines mark statistical thresholds (FDR ≤0.05 and absolute value of the log2 FC ≥1). **(C)** Representative fusion transcript isoform structures from the *ZNF254*::*GNAS* reported GF in the 5’-3’ direction, with gene strandness denoted in the title. The reported fusion breakpoints (red vertical lines) connect a novel *ZNF254* isoform (NIC, “novel in catalog,” lacking expression quantification) to an annotated *GNAS* transcript (GNAS-238), with base pair resolution expression shown as a color gradient (white to purple), transcript exon positions from IsoQuant reported isoform structures. In *ZNF254*, the breakpoint occurs downstream of exon 3, resulting in a novel terminal exon, while in the *GNAS* gene the breakpoint occurs upstream of exon 2 relative to the MANE Select reference transcript (dark blue). Color intensity represents exon-level expression, with matching exons in blue (*ZNF254*) or purple (*GNAS*) and non-matching exons in gray.

In parallel, we observed an extensive landscape of novel (previously unannotated) isoforms in glioma. IsoQuant detected 132,716 novel transcripts (65,580 novel-in-catalog and 67,136 novel-not-in-catalog) from 12,380 protein-coding genes, while FLAIR identified 143,938 novel isoforms from 11,228 protein-coding genes. Novel isoforms were abundant in both HGG and LGG and often clustered within specific genes, suggesting isoform-switching events that could alter protein structure or regulation. Several genes, including *FAM228B*, *PAQR6*, *LST1*, *TRPT1*, *BCAN*, and *NDRG2*, produced multiple distinct novel isoforms detected by both tools and showed substantial expression in external glioma RNA-seq data from the Human Protein Atlas and TCGA, with *BCAN* and *NDRG2* displaying cancer-enhanced expression in glioma.^36^ These findings highlight the transcriptomic diversity and prevalence of previously unannotated isoforms in glioma and provide a foundation for linking novel transcript structures to genes involved in GFs.

We examined transcript isoforms overlapping annotated fusion breakpoints to investigate the relationship between isoform diversity and GFs. Given the low overlap between significant DE isoforms identified by IsoQuant and FLAIR, this analysis considered the union of known and novel isoforms detected by either tool, rather than restricting to consensus isoforms, to capture the full diversity of transcript structures revealed by long-read sequencing. We focused on fusion partner genes in which at least one transcript (known or novel) exhibited non-canonical transcript start or end positions proximal to the reported fusion breakpoint in the same glioma sample. A subset of the fusion-related isoforms exhibited fusion-like structural features, such as premature transcript termination sites within the 5’ gene and novel transcriptional initiation sites within the 3’ gene partner. These features were observed at genomic regions consistent with a subset of the reported fusion breakpoints, reflecting positional overlap between transcript boundaries and fusion breakpoints. As a representative example, the *ZNF254*::*GNAS* GF (**Figure 2C**) showed a novel *ZNF254* isoform classified as novel-in-catalog that terminated downstream of exon 3, while the corresponding 3’ gene partner aligned to an annotated *GNAS* transcript initiating at exon 2. Additionally, the combination of a lowly expressed, novel upstream transcript (*ZNF254*) with a highly expressed, downstream transcript (*GNAS*) follows a pattern consistent with fusion-like structures. Similar breakpoint-proximal isoform partners were observed for additional fusions, such as *ATG4B*::*CNTRL*, *ACVR1B*::*CCDC60*, *PTPRK*::*NOX3*, and *DUSP22*::*APOE*. Together, these results illustrate how long-read isoform profiling complements GF detection by providing transcript-level context near reported fusion breakpoints and highlighting candidate fusion events.

### Functional Validation of Selected GFs in a Drosophila Tumor Testing Model

We curated a subset of GFs with breakpoints located within CDS/near exon boundaries to prioritize candidate fusions for experimental validation. These fusions have breakpoint configurations that are more likely to generate in-frame, functionally relevant fusion transcripts suitable for downstream functional validation in *Drosophila*. In our previous work, 20 GFs were validated computationally by visualizing supporting reads and experimentally by PCR and Sanger sequencing. From the additional 25 HGG and LGG samples presented here, we evaluated five additional fusions, *ABL1*::*WDR83OS*, *BCL11A*::*VIRMA*, *PRICKLE1*::*FHIT*, *MDM2*::*NUP107*, and *DDX6*::*DCTN2*. Long-read alignments supporting each fusion visualized in IGV and Genome Ribbon, (**Supplemental Figure S1A**) and experimental PCR amplification across fusion junctions (**Supplemental Figure S1B**) confirmed these GFs. Of these, *ABL1*::*WDR83OS* and *MDM2*::*NUP107* were further validated with Sanger sequencing chromatograms (**Supplemental Figure S1C**), providing orthogonal evidence that these GFs represent true events rather than technical artifacts. Across the validated fusion cohort identified from all 49 glioma samples we selected 15 high-confidence GFs for functional testing in *Drosophila* and generated expression constructs using breakpoint-defined fusion sequences, as summarized in **Table 1**. These constructs were designed to test the functional consequences of precise fusion breakpoints rather than the broader isoform landscape observed by long-read sequencing. Of these 15 GFs, three included at least one gene from CHOP FUSIP and 14 included at least one gene from the COSMIC Cancer Gene Census, supporting the clinical and biological relevance of the GF set (**Table 1**).

**Table 1.**
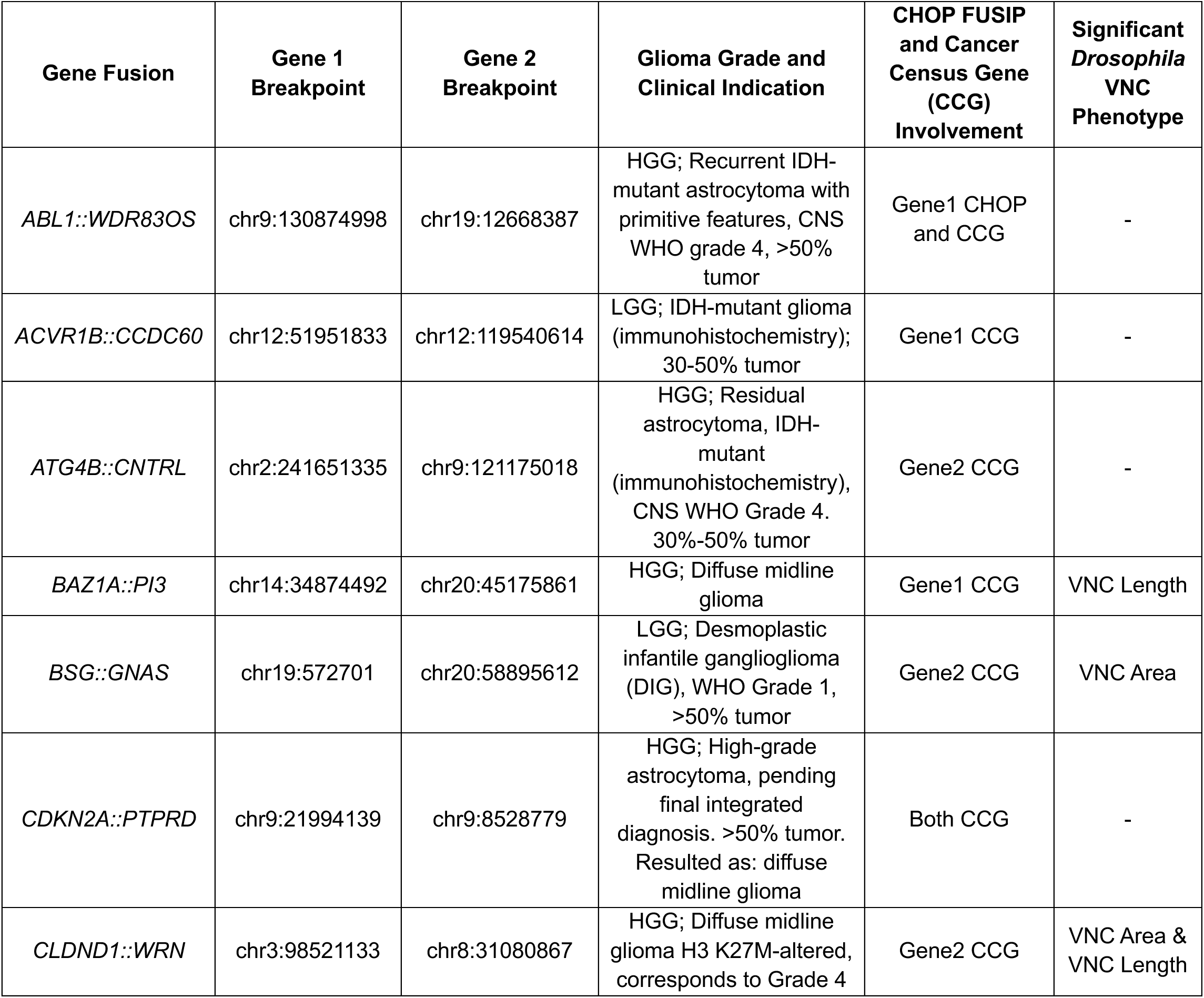
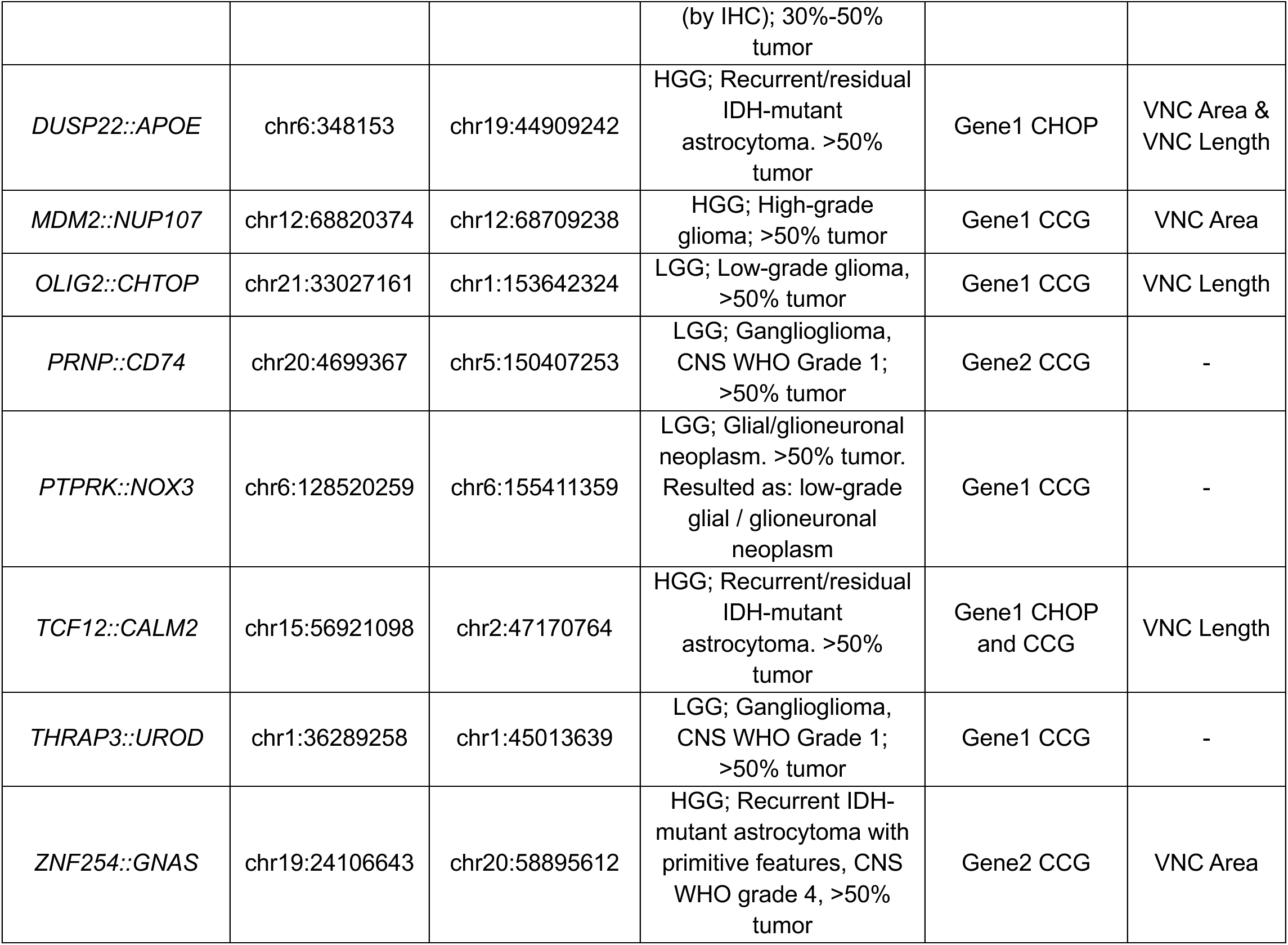
Summary of prioritized GFs induced in *Drosophila* tumor model and resulting significant oncogenic phenotypes. Summarized novel GFs with their breakpoints, glioma grade, clinical indications, cancer census gene (CCG) and CHOP FUSIP involvement, and significant *Drosophila* validation results. GF breakpoints were refined based on Sanger sequencing base-pair resolution confirmation. The glioma grade and clinical indication reflect the diagnostic and histopathologic context of the patient sample. CHOP FUSIP and Cancer Census Gene (CCG) involvement denotes whether either fusion gene partner is a known cancer gene or included on the clinical CHOP FUSIP. Significant *Drosophila* VNC phenotype indicates if the GF induced significant measurable changes in VNC length or area, and the dash indicates no significant phenotype.

To assess *in vivo* effects, we generated transgenic *Drosophila* lines using the pan-glial *repo-GAL4*^37^ driver to express an *mRFP*^38^ reporter to label glia and to overexpress each fusion gene transcript (**Supplemental Table S6**). We focused on phenotypes previously associated with oncogenic signaling including tumor-like overgrowth and altered glial cell proliferation or migration.^29^ Previously, we established five pediatric LGG *Drosophila* models using patient-derived GFs (*KIAA1549::BRAF, QKI::NTRK2, FGFR::TAAC1, QKI::RAF1, MKRN::BRAF*) and demonstrated that individual GFs produced distinct, quantifiable phenotypes such as VNC area, length, and glial migration (**Figure 3**).^32^ Consistent with this framework, multiple GFs in the current study induced clear morphological abnormalities in the VNC compared to wild-type (WT) controls as shown in **Figure 4**. In this model, an increase in VNC area reflects excessive glial growth, whereas VNC elongation may indicate developmental defects in tissue organization. Although these phenotypic measurements do not distinguish among specific cellular mechanisms, they provide a useful *in vivo* readout of fusion-induced disruption of glial cell regulation. Visually, *CLDND1*::*WRN* and *DUSP22*::*APOE* produced VNC elongation and thickening, whereas *OLIG2*::*CHTOP* and *TCF12*::*CALM2* resulted in more elongated and distorted VNC structures (**Figure 4A**). Quantitative analysis of VNC area (**Figure 4B**) and length (**Figure 4C**) revealed that 8 GFs induced significant changes (p ≤0.05) in one or both measurements relative to WT. Notably, *CLDND1*::*WRN* and *DUSP22*::*APOE* exhibited some of the strongest elongation phenotypes, with markedly increased VNC length and area comparable to established glioma-associated fusions such as *QKI*::*NTRK2*, *FGFR1*::*TACC1*, and *KIAA1549::BRAF*.^32,39,40^ These phenotypes are consistent with conserved mechanisms of aberrant glial proliferation and patterning driven by the breakpoint-defined glioma fusions.

**Figure 3.**
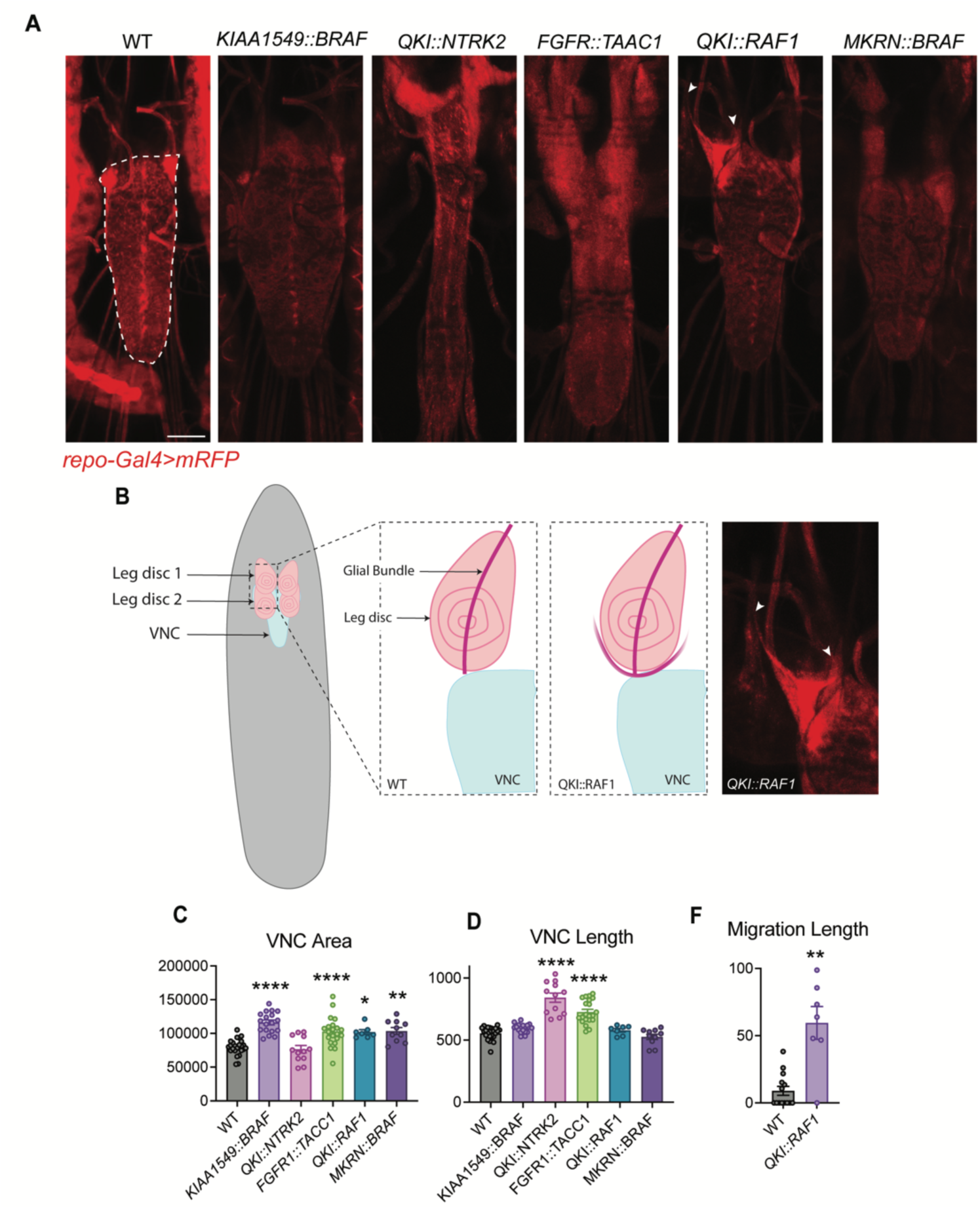
Patient-derived glioma GFs induce quantifiable VNC phenotypes in *Drosophila* models of LGGs. **(A)** *In vivo* images of third instar larval VNCs, with *repo-GAL4* driving *mRFP* expression and GFs in glia. A white dashed line outlines the WT VNC. Scale bar = 200 µm. **(B)** Schematic illustrating typical glial migration patterns in the leg imaginal discs of third instar WT and *QKI::RAF1* overexpressing larvae. *In vivo* image expressing mRFP in glia in *QKI::RAF1* overexpressing larvae, white arrows indicate aberrant migratory glial cells. **(C)** Quantified VNC area (μm^2^), **(D)** VNC length (μm), **(E)** and glial cell migration distance (μm). Data are mean ± SEM, analyzed by one-way ANOVA followed by Dunnett’s multiple comparisons test, *P<0.05, **P<0.01, ***P<0.001, ****P<0.0001. N > 7 larvae.

**Figure 4.**
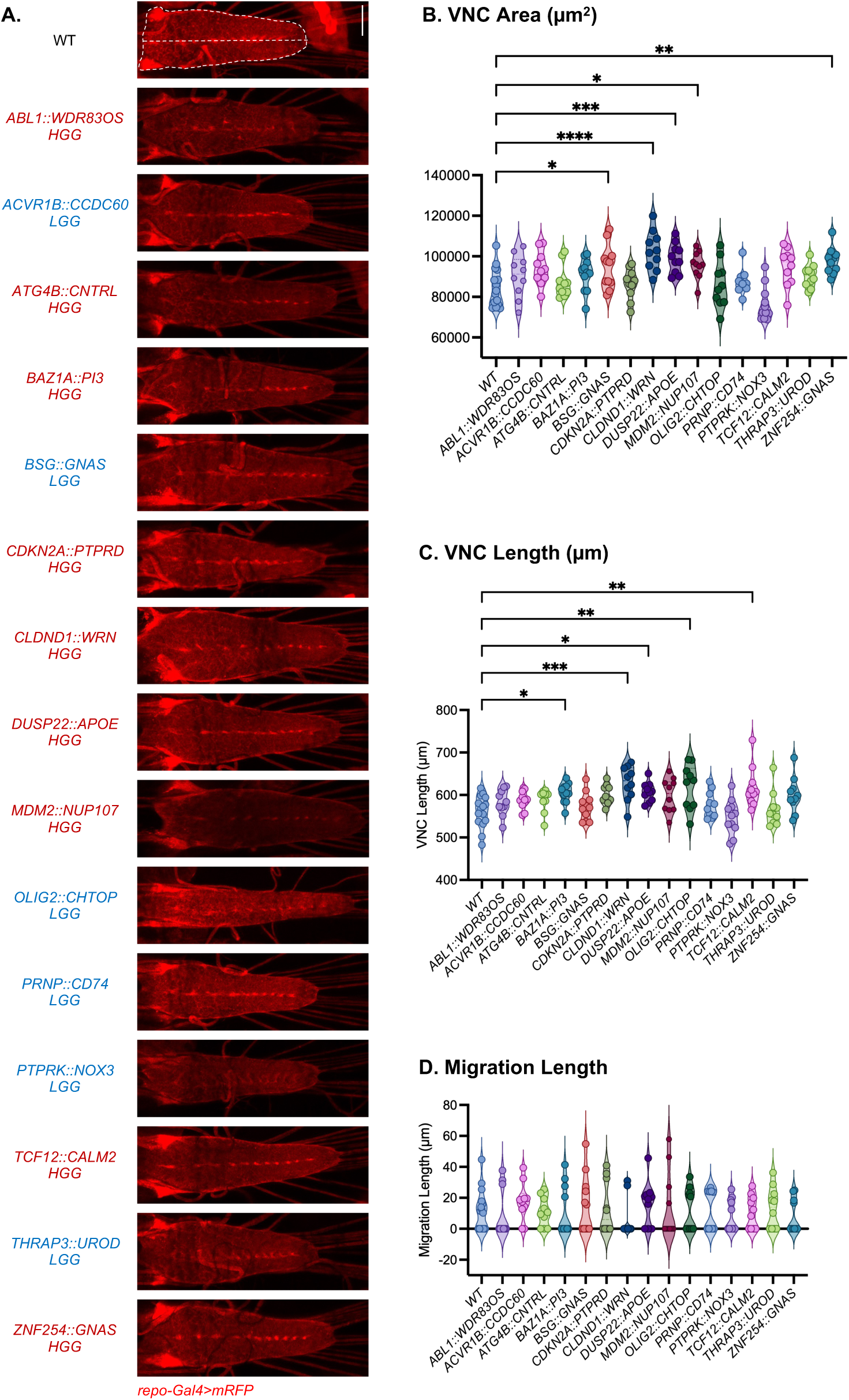
Phenotypic characterization and quantification of glial overexpression of breakpoint-defined GFs in *Drosophila melanogaster* transgenic lines. **(A)** *In vivo* images of third instar larval VNCs with *repo-GAL4* driving *mRFP* expression and GFs in glia. Fusion names are labeled and color-coded by tumor grade: HGGs are red and LGGs are blue. Scale bar = 200 µm. **(B-D)** Quantification of VNC morphology and glial cell migration phenotypes. **(B)** VNC area (μm^2^). **(C)** VNC length (μm). **(D)** Glial cell migration distance (μm) along leg imaginal disc edges. Data are mean ± SEM, analyzed by one-way ANOVA followed by Dunnett’s multiple comparisons test, *P<0.05, **P<0.01, ***P<0.001, ****P<0.0001. N > 10 larvae.

Additionally, we quantified glial migration by measuring migration distance along the VNC (**Figure 4D**). This analysis was motivated by our previous work demonstrating that glial overexpression of the GF *QKI::RAF1* in *Drosophila* produced a pronounced infiltrative phenotype, characterized by aberrant glial migration, and models the diffuse nature of pediatric-type diffuse LGGs. The aberrant glial migration is characterized by glia migrating beyond the CNS and into leg imaginal discs, the epithelial structures adjacent to the CNS. This migration defect reflects the infiltrative, diffuse nature of pediatric LGGs, highlighting the utility of glial migration as a functional readout of tumor infiltration (**Figure 3B**). In this cohort of GFs we observed that several fusions altered migration relative to WT controls, although none reached statistical significance. Combined with structural changes including VNC area and length, these migration defects support a model in which specific glioma GFs perturb multiple aspects of the nervous system architecture.

Overall, 8 of the 15 breakpoint-defined GFs (∼53%) induced significant VNC alterations in at least one VNC phenotype (area or length), indicating a substantial fraction of long-read prioritized GF candidates exhibited measurable oncogenic-like activity *in vivo*. These findings demonstrate that GFs outside of targeted fusion panels can encode proteins capable of driving tumor-like overgrowth and structural disruption *in vivo*, highlighting their oncogenic potential, and supporting the integration of additional long-read fusion profiling as a complementary approach in challenging or FUSIP-negative glioma cases.

In contrast, seven fusions, including *ABL1*::*WDR83OS*, *ACVR1B*::*CCDC60*, *ATG4B*::*CNTRL*, *CDKN2A*::*PTPRD*, *PRNP*::*CD74*, *PTPRK*::*NOX3*, and *THRAP3*::*UROD*, did not differ significantly from WT in any of the measured phenotypes. To confirm that the absence of observable phenotypes was not due to insufficient transgene expression, we assessed fusion construct expression levels by immunostaining of the FLAG tag. We observed robust expression across these lines, with the FLAG signal detected in both glial processes and nuclei, as shown in **Supplemental Figure S3**. Together, these results demonstrate that a subset of breakpoint-defined glioma GFs is sufficient to drive tumor-like VNC overgrowth, elongation, and structural disruption *in vivo*, whereas other fusions may require additional genetic or cellular alterations to exert detectable biological effects.^29^

### Pathway Enrichment Analysis

Pathway enrichment analysis was performed on four distinct gene sets to characterize the biological processes disrupted in glioma: (1) genes involved in detected GFs, (2) genes with significantly DE isoforms (consensus between IsoQuant and FLAIR), (3) genes with novel isoforms from both IsoQuant and FLAIR, and (4) genes involved in GFs that produced significant phenotypic alterations in *Drosophila*. Gene set enrichment was assessed using Enrichr with the KEGG 2021 Human and Elsevier Pathway Collection databases, and we then examined the significantly enriched pathways for glioma- and CNS-related terms (see Methods).

Genes involved in GFs showed enrichment for broad cancer-related processes and multiple tumor types. In KEGG, significant pathways included “pathways in cancer” and cancer types such as chronic myeloid leukemia, colorectal cancer, non-small cell lung cancer, melanoma, and glioma, in which driver GFs have been identified. Elsevier highlighted genes and proteins in pathways associated with altered expression in cancer, “including cancer-associated proliferative signaling”, “glioblastoma (secondary and primary)”, “astrocytoma”, and “proteins involved in both glioblastoma and astrocytoma”. Fusions in which both breakpoints were in CDS or near-exon regions were most significantly enriched for “pathways in cancer” and “transcriptional misregulation in cancer,” consistent with a subset of events most likely to result in biologically functional or oncogenic fusion transcripts. Of these genes, pathways such as “proteins with altered expression in cancer metastases” was the most significant, and “proteins involved in astrocytoma and glioblastoma”, were among the top 15 significant pathways, across all comparisons between glioma samples and controls.

Genes with significant DE isoforms showed more consistent and pronounced enrichment for glioma- and CNS-related pathways than the fusion genes. In KEGG, consensus DE isoform genes were enriched for neurological diseases and neurodegeneration pathways, including “Alzheimer’s disease”, “Parkinson disease”, “Amyotrophic Lateral Sclerosis (ALS)”, “pathways of neurodegeneration,” as well as “glioma.” Enrichment using the Elsevier Pathway Collection likewise identified glioma-related terms such as “proteins involved in glioma,” “glioblastoma,” and “astrocytoma,” alongside “neuroblastoma- and ALS-related signaling”. Among significantly DE genes from the HGG vs LGG comparison, genes were strongly enriched for mitotic and cell-cycle pathways, including “metaphase/anaphase phase transition”, “spindle assembly”, and “APC/C-CDC20 complex”, indicating that isoform-level differences between high- and low-grade gliomas were concentrated in cell division and the chromosome-level.

Novel isoform genes followed a similar overall pattern. Using the union of genes with novel isoforms from IsoQuant and FLAIR, KEGG enrichment highlighted pathways related to neurodegeneration and neuronal function (ALS, Huntington’s disease, Parkinson’s disease, and pathways of neurodegeneration) as well as cellular processes such as protein processing in the endoplasmic reticulum, endocytosis, and cell cycle. Elsevier pathways for novel isoform genes included “proteins involved in ALS” and “cell cycle-related modules”. The proportion of glioma/CNS-related pathways among significant hits was comparable to those observed for genes with DE isoforms, suggesting that novel isoforms frequently arise in genes involved within glioma-relevant signaling networks. Collectively, these pathway analyses indicate that both fusion-associated and isoform-level alterations contribute to overlapping glioma and CNS pathways, reinforcing that structural rearrangements and transcriptomic dysregulation act together to shape glioma biology.

To further contextualize GFs that exhibited significant phenotypic alterations in the *Drosophila* functional validation assays, pathway enrichment analysis identified pathways directly relevant to glioma biology. Despite the small size of this gene set, 15 genes involved in 8 fusions, the most significant KEGG glioma-associated pathway was “glioma,” while enrichment using the Elsevier Pathway Collection highlighted categories including “proteins involved in astrocytoma”, both among the top 15 significant pathways. Additional significant glioma-related pathways from Elsevier included “glioblastoma, proneural subtype”, “proteins involved in glioma”, “non-hereditary genetic rearrangements in neuroblastoma”, and “astrocytoma”, similar to the enriched pathways observed for genes with significant DE isoforms. In addition, neurodegeneration-related pathways, including Alzheimer’s and Huntington’s disease were also enriched, consistent with shared molecular and cellular mechanisms between neurodegeneration and glial dysfunction. Together, these results indicate that GFs capable of driving observable *in vivo* phenotypes preferentially involve genes involved in oncogenic and CNS-associated signaling networks, supporting the biological relevance of fusions prioritized through functional validation.

## METHODS

### Clinical Cohort and Sample Characteristics

This study analyzed a cohort of 49 glioma cancer patient samples, including 24 in our previous study on combining panel-based and whole-transcriptome-based gene fusion detection^24^ and an additional 25 samples processed using identical protocols. We only used whole transcriptome data in the current study. The final cohort consisted of 24 low-grade gliomas (LGG) and 25 high-grade gliomas (HGG) distributed across both sample sets. All samples were obtained from the Division of Genomic Diagnostics (DGD) at the Children’s Hospital of Philadelphia (CHOP), under approved protocols, and de-identified prior to analysis. Clinical indications from the DGD at CHOP for all 49 samples, including the 25 additional samples presented here, are detailed in **Supplemental Table S1**. Prior to analysis, the DGD at CHOP comprehensively characterized all 49 samples for disease-causal gene fusions (GFs) using the CHOP Cancer Fusion Panel.^22^ This fusion panel utilized short-read sequencing on the Illumina platform, which targets over 700 exons from 117 cancer genes (updated to 119 genes) known to be involved in GFs. All samples were deemed fusion panel-negative for well-known disease-causal GFs. The samples were later subjected to Oxford Nanopore Technologies (ONT) long-read whole-transcriptome sequencing for comprehensive GF and transcript isoform analysis, with downstream computational and experimental validations of high-confidence potentially novel GFs.

### Long-read Whole-Transcriptome Sequencing

All 49 samples were sequenced using long-read whole-transcriptome sequencing on the ONT platform. Library preparation methods for the 24 previously analyzed samples utilized cDNA-PCR sequencing, as detailed in our previous publication.^24^ For the remaining 25 samples, two distinct ONT library preparation methods were employed across three sequencing pools, numbered 1, 4, and 6. Pool #1 (samples #33, 35, 36, 37, 41, 46, 47, 50, 60, and 67) libraries were prepared using ONT’s Direct cDNA Sequencing protocol (DSC_9187_v114_revH_19Apr2023) where cDNA was first synthesized from total RNA. This was followed by cDNA amplification using the Low Input by PCR protocol (LWP_9183v114_rev_07Mar2023), after which the amplified cDNA was barcoded and sequenced using the Native Barcoding Kit 24 (V14) (SQK-NBD114.24). While the recommended input for cDNA synthesis is 1 μg of total RNA, 250 ng of total RNA was used instead. All other steps in the protocols were followed without further modifications. Pools #4 and #6 (remaining samples) libraries were prepared using the cDNA-PCR sequencing method, which involved ONT’s cDNA-PCR Sequencing V14 Barcoding Kit (SQK-PCB114.24) protocol. K562 cell line samples were included as controls in both pools (#4 & #6). 250 ng of input total RNA was used instead of the recommended 500 ng and the beads ratio in the final cleanup step was adjusted from 0.7x to 0.4x. All remaining steps in the protocol were followed without further modifications.

Following ONT library preparation, samples were multiplexed on individual R10.4.1 PromethION flow cells and sequenced for 72 hours using our in-house P2 Solo sequencer. Each sequencing pool contained a mixture of high-grade glioma (HGG) and low-grade glioma (LGG) samples, except for pool 6, which included only one HGG (a resequencing of sample #32) and seven LGG samples. The distribution of samples across the sequencing pools included: pool 1 (7 HGG, 3 LGG), pool 2 (5 HGG, 3 LGG), pool 3 (3 HGG, 4 LGG), pool 4 (3 HGG, 5 LGG), pool 5 (6 HGG, 2 LGG), and pool 6 (1 HGG, 7 LGG). After sequencing, internal basecalling was performed using ONT’s software. **Supplemental Table S2** summarizes the sample input concentrations and sequencing data generated, and **Supplemental Table S3** provides detailed sequencing generation metrics at various timepoints for the PromethION flow cell sequencing.

### Gene Fusion Detection and Analysis

Comprehensive GF detection was performed on all 49 samples using our previously established long-read whole-transcriptome GF detection pipeline.^24^ The pipeline employs an ensemble-based approach utilizing three complementary GF detection tools including, LongGF^41^, JAFFAL^42^, and FusionSeeker^43^. Fusion calls from each tool were combined and duplicates were removed in the following steps: (1) removal of exact duplicates from the same GF program (identical genes and breakpoints) and those within a 50 bp breakpoint window for both fusion partners across GF programs, and (2) identification of reciprocal fusions (same gene partners in reversed orientation within 50 bp of respective breakpoints). For reciprocal pairs, the orientation with higher read support was retained and both fusion entries were kept if read support was equal. Standard filtering parameters from the GF detection pipeline were then applied, including, fusions were retained if at least one gene was from the CHOP Fusion Panel or Cancer Census Gene lists, minimum 3 supporting reads, removal of pseudogenes, ribosomal proteins, HLA genes, and mitochondrial fusions, and exclusion of fusions with inconsistent strand orientation. This relatively low read-support threshold was intentionally selected to maximize sensitivity in a cohort that had initially tested negative by a targeted fusion panel and therefore was expected to have rare, non-recurrent, or lowly expressed fusion transcripts. To focus the analysis on brain-related fusions, we excluded fusions identified in healthy brain tissue samples from the Genotype-Tissue Expression (GTEx) long-read RNA-seq project (v9)^44^. This combined ensemble detection and filtering strategy enhanced identification of both known and potentially novel GFs that may have been overlooked by short-read sequencing.

For breakpoint location type annotation, we compared both GENCODE (v45)^45^ MANE Select and the longest available protein-coding transcript isoforms for each gene. In cases where a breakpoint was classified as intronic by one but exonic by the other, breakpoint location types were reclassified as exonic. We prioritized the longest transcript as it may better capture isoform diversity. Breakpoints were then classified as exonic, intronic, intergenic, or within untranslated regions (UTRs). Intronic breakpoints located within 200 bp of an exon boundary were additionally reclassified as exonic to account for potential alignment ambiguity at exon-intron junctions or alternative splicing events. This 200 bp window was chosen because most fusion junctions fall at or very close to exon-exon boundaries, and long-read fusion tools explicitly use exon boundaries to define high-confidence breakpoints.^42^ Intronic splicing regulatory elements are also concentrated within ∼100 bp of splice sites, so a 200 bp margin gives a conservative buffer for long-read mapping while still restricting analysis to splice-proximal breakpoints that are more likely to generate expressed chimeric transcripts.^32^ GFs with both breakpoints classified as intronic, intergenic, or unannotated were removed from downstream analyses.

### Gene Lists for Fusion Annotation and Prioritization

To systematically prioritize clinically relevant GFs in the previously fusion panel-negative glioma samples, we developed a comprehensive annotation strategy using multiple cancer- and glioma-specific curated gene lists. For each gene identified in the fusions, we first annotated for involvement in the list of 119 targeted CHOP fusion panel genes to identify fusions potentially overlooked by the panel initially with established clinical relevance. Next, oncogene and tumor suppressor gene(TSG) classifications were recorded using the COSMIC Cancer Gene Census^46^, OncoKB^47^, ONGene^48^, and TSGene 2.0^49^ databases to provide a comprehensive functional characterization of the genes and its role in the cancer.

We further annotated genes based on both the general COSMIC Cancer Gene Census and a curated glioma-specific subset, which we generated by extracting genes associated with glioma or CNS tumors. This was accomplished by searching the tumor type annotations for glioma-related keywords (glioma, astrocytoma, glioblastoma, oligodendroglioma, and CNS) and excluding irrelevant types such as paraganglioma.

To assess gene essentiality in glioma, we utilized the Sanger Cancer Dependency Map (DepMap)^50,51^, a large-scale CRISPR-Cas9 screening resource that identified genes essential for cancer cell survival and proliferation across more than a thousand cancer cell lines. We filtered the DepMap cell lines for those associated with CNS and brain, focusing on those with “Diffuse Glioma” as the primary disease and including all subtypes (glioblastoma, astrocytoma, anaplastic astrocytoma, oligodendroglioma, and gliosarcoma). For each glioma subtype-related cancer cell line, maximum dependency probabilities were calculated for each gene, and those with scores 0.9 or above were classified as highly essential candidates. Correspondingly, minimum gene effect scores were calculated, and genes with scores -1 or less were identified as potential glioma-essential genes. The lenient thresholds were used to capture a broad range of glioma-related genes and ensure comprehensive prioritization in downstream analyses.

Additionally, genes were annotated for their involvement in reported GFs from brain and CNS-related diseases using the Mitelman Database^26^, as previously described in our long-read whole-transcriptome GF detection pipeline. This integrated approach enabled comprehensive annotation of genes in the fusions with potential clinical relevance and functional importance to guide glioma-focused downstream analyses.

### Transcript Isoform Detection and Differential Expression Analysis

All 49 glioma and 22 long-read sequenced control brain tissue samples, sourced from GTEx (v9), were analyzed using both IsoQuant^52^ and FLAIR^53^ isoform detection tools, each run with default parameters, GENCODE (v46) gene annotation, and GRCh38 reference genome. For FLAIR, we optimized the isoform detection by splitting reads from each sample into two groups based on chromosome (chromosomes 1-10 and the remaining chromosomes). Isoform detection was performed separately on each group, and all detected transcripts were subsequently combined prior to FLAIR’s transcript collapse step, which merged redundant and structurally similar transcripts to produce a set of nonredundant isoforms for downstream analysis. IsoQuant isoform detection did not require this processing modification.

Detected transcripts from both tools were initially filtered to retain only those present in the reference annotation. For IsoQuant, transcripts were further required to be uniquely assigned to a single isoform to exclude reads that could not be distinguished between multiple possible isoform candidates. From both IsoQuant and FLAIR analyses only protein-coding transcripts, excluding those on the mitochondrial chromosome, were retained. For each sample, transcript-level expression was quantified by both raw read counts and transcripts per million (TPM), with undetected transcripts assigned a value of zero. For each transcript, the average expression values and percentile ranks were calculated across all glioma and control samples, and between HGG and LGG glioma subgroups.

#### Differential Transcript Isoform Expression

To identify differentially expressed isoforms, edgeR^54^ (v4.4.2) and DESeq2^55^ (v1.46.0) were applied in R (v4.4.3). Filtered isoform count matrices from IsoQuant and FLAIR were analyzed across four comparisons: all gliomas vs. controls, HGG vs. controls, LGG vs. controls, and HGG vs. LGG. Expression-based filtering was applied to remove noisy or lowly expressed isoforms, where transcripts were first required to have ≥10 counts in a minimum number of samples equal to the smallest group size in each comparison and ≥15 total counts for a given isoform when summed across all samples in the comparison. For the HGG vs LGG comparison, the sequencing pool was included as a covariate to correct for potential batch effects. This covariate was not applied to the other comparisons as the control sequencing pool information was not available.

Differential expression was assessed using the quasi-likelihood F-test in edgeR and the Wald test in DESeq2. Isoforms were considered significantly differentially expressed (DE) if the false discovery rate (FDR) in edgeR or the adjusted p-value in DESeq2 was ≤0.05 and the absolute value of the log2 fold change (FC) in expression was ≥1. Results from each method were ranked by significance and cross-referenced at the gene level with genes involved in identified GFs. To generate high-confidence consensus DE isoforms, significantly DE isoforms with concordant fold-change direction (both upregulated or both downregulated) from both edgeR and DESeq2 were retained. To summarize the results across tools, we calculated the mean log_2_ fold change and the maximum FDR value between the two methods to represent statistical significance.

#### Novel Transcript Isoform Detection

IsoQuant and FLAIR were used to identify novel transcript isoforms, defined as isoforms absent from the reference GENCODE annotation, in glioma patient samples. IsoQuant classified novel isoforms as either “novel in catalog” (NIC) or “novel not in catalog” (NNIC) and provided exon interval information with TPM values for quantification. FLAIR also provided exon interval information for novel isoforms but lacked read support for individual transcripts, preventing accurate TPM estimation. Novel isoforms detected from both tools were filtered to exclude mitochondrial genes and retain only those from protein-coding genes.

To identify recurrent novel transcript structures, a multi-step grouping strategy was applied to the filtered novel isoforms from both tools. First, transcripts were first grouped by the total number of exons. Within each group, transcripts were further subdivided based on transcript start position similarity (within 100bps). For each subgroup, exon start and end positions were iteratively compared, and isoforms were grouped together if all corresponding exon boundaries fell within 100bps of each other. The average exon coordinates within each isoform group served as the representative structure for that group. To identify the most abundant novel isoform structure per gene, the group containing the largest number of similar isoforms was selected. Finally, genes were ranked by the total number of distinct novel isoform structures to highlight genes with recurrent novel isoform patterns across glioma samples.

#### Overlap of GF Breakpoints with Differentially Expressed and Novel Transcript Isoforms

To investigate the relationship between the detected GFs and transcript isoforms, we systematically identified all fusions that contained at least one gene with at least one significantly differentially expressed isoform based on gene name. For isoforms identified in these fusion genes, transcript structures were analyzed to determine whether their non-canonical stop and new start sites were located proximal to the fusion breakpoints in the corresponding samples where fusions were detected. Specifically, for each gene in a fusion, we evaluated whether at least one transcript (known or novel) had a non-canonical transcript start or end position located within 10% of the gene length from the reported fusion breakpoint, with a minimum distance of 500 bp and a maximum of 2 kb. Thresholds chosen to account for gene length variability while restricting to breakpoint-proximal transcript changes. This proportional threshold was chosen to account for gene length variability. Non-canonical stops in the first fusion partner gene and novel starts in the second fusion partner gene were examined for alignment with the respective transcript structures.

All transcript isoforms that overlapped GF genes and near fusion breakpoints were visualized to examine the isoform structures and expression patterns in both fusion partner genes. Exon boundaries of transcripts were defined as the average of all detected exon start and end coordinates across supporting reads. Exon-level read counts from IsoQuant were used to generate expression profiles across each isoform, allowing visualization of relative expression changes along isoform and exon structures. For isoforms detected only by FLAIR, transcript structures were visualized, but expression profiles could not be generated as FLAIR did not retain exon-level read support. When multiple transcripts were present, isoforms were ranked by expression level (TPM) to prioritize the most abundant structures. Importantly, these isoform analyses were used to provide transcript-level context for genes involved in fusion breakpoints and were not used to design the *Drosophila* fusion constructs themselves. The *in vivo* constructs were based on patient-specific breakpoint coordinates mapped onto reference transcripts.

### Experimental Validation and Functional Characterization in Drosophila

#### Validation of Potentially Novel GFs

Potentially novel GFs identified within our dataset were initially validated computationally using the Integrative Genomics Viewer (IGV) and Genome Ribbon (**Supplemental Figure S1A**). These GFs were further experimentally validated by PCR amplification using specifically designed primers (**Supplemental Table S7**) around the reported fusion breakpoints. Amplified PCR products were confirmed by gel electrophoresis of the expected product size and subjected to Sanger sequencing to verify the fusion breakpoint present in the samples (**Supplemental Figure S1B-C**).

#### GF Drosophila Construct Design

Fusion cDNA sequences for *Drosophila* transgenic expression were designed based on Sanger-validated fusion sequences. GENCODE transcripts with coding sequence (CDS) or near exon (<200 bps) were prioritized as the reference sequence of each gene, where the reported fusion breakpoints were adjusted to match the experimentally confirmed fusion Sanger sequences where possible. Open reading frame (ORF) analysis was performed on all fusion sequences to identify the presence of premature stop codons. Fusion sequences with both gene sequences in-frame with the original gene’s frame and a complete ORF were prioritized for *Drosophila* expression. For fusions with genes out of frame from the original gene’s frame and containing premature stop codons were also included for expression in *Drosophila*, using the sequence upstream the premature stop codon to mimic likely translation of the functional fusion protein product. To account for variability in patient fusion isoforms, fusion transcripts were designed to express breakpoint-defined ORFs rather than full-length fusion transcripts. All fusion sequences, breakpoint coordinates, transcript IDs, and ORF sequences used for *Drosophila* expression are provided in **Supplemental Table S6**.

The *BCL11A*::*VIRMA* fusion had a computationally annotated *BCL11A* breakpoint located within the 5’ UTR region (ENST00000642384.2). Sanger sequencing confirmed the 5’ UTR annotation in the patient sample, validating our computational pipeline. However, since UTR regions are not translated and have minimal contribution to fusion protein function, the resulting *Drosophila* fusion construct was designed with the entire *BCL11A* coding sequence, fused to the sequence downstream of the *VIRMA* breakpoint. Another exception included the *ABL1*::*WDR83OS* GF, as the fusion construct was generated using the predicted breakpoints. The Sanger sequencing later revealed a 7-nucleotide microhomology region at the fusion junction that was present in both gene reference sequences but was not assigned to either gene from the GF detection program (JAFFAL). This resulted in a 415 amino acid fusion sequence with a premature stop codon, whereas Sanger confirmed the fusion would result in a 417 amino acid fusion with two additional residues and no premature stop codon. The model construct retains the complete *ABL1* protein including all functional domains (SH3, SH2, protein kinase domain; InterPro), while the *WDR83OS* portion contributes only a small sequence that does not contain annotated functional protein domains. The 2-amino acid difference occurs exclusively within the end of the *WDR83OS* non-functional gene region.

#### Generation of Drosophila Fusion Transgenic Lines

The synthesized fusion cDNA transcripts were ordered from GenScript. The fusion coding sequences were cloned into the pACU2 fly expression vector, with an N-terminal FLAG tag. The plasmids were then injected into fly embryos via φC31-mediated site-specific insertion (Rainbow Transgenic Flies Inc) to generate transgenic fly strains. All fly crosses, including larvae, were kept at 25 °C. Live imaging was conducted as previously described.^56,57^ Briefly, randomly selected female and male third instar larvae 5-6 days after egg lay (AEL) were collected in vials containing standard fly food and screened for the correct genotype. Larvae were anesthetized using diethyl ether and individually mounted in 90% glycerol under coverslips sealed with grease. The Zeiss LSM 880 microscope was used for live imaging. To quantify the infiltrative phenotype, the length that glial cells traveled along the edges of the leg disc (in μm) was measured using the FIJI software. *Repo-Gal4*^37^ and *UAS-mRFP*^38^ have been previously described.

#### Immunohistochemistry

Third instar larval VNCs 5-6 days AEL were dissected in phosphate-buffered saline (PBS) and fixed for 20 minutes in 4% paraformaldehyde. Samples were washed 3x in 0.3% PBS-T (PBS + Triton X), blocked for 1 hour in 5% normal donkey serum, and incubated overnight at 4°C in a primary antibody solution consisting of 1:500 mouse anti-FLAG antibody (Sigma F3165) diluted in blocking buffer. Following incubation, the primary antibody was washed 3x with PBS-T and incubated for 2 hours at room temperature in 1:200 donkey anti-mouse Alexa Fluor 488. Samples were then washed 3x in PBS-T, incubated for 20 minutes in a 1:1 PBS-T:glycerol solution before mounting in Vectashield to be imaged with a Zeiss LSM 880 confocal microscope and analyzed using FIJI software.

### Pathway and Functional Enrichment Analysis

To identify biological pathways associated with genes involved in GFs and significantly DE isoforms, we performed gene-level over-representation analysis using Enrichr gene-set libraries, accessed via the GSEApy v1.1.10 Python package.^58–61^ Pathway enrichment was performed separately for multiple gene sets, including (1) genes involved in detected GFs, (2) genes with significantly DE isoforms identified by IsoQuant and FLAIR, (3) genes with novel isoforms identified by either tool, and (4) genes involved in GFs that produced significant phenotypic alterations in *Drosophila*. Across all glioma comparisons, unique genes from each set were used as input for enrichment analysis.

Enrichment was evaluated across curated gene-set libraries, including the KEGG 2021 Human and Elsevier Pathway Collection. Statistical significance was assessed using Enrichr adjusted P values, and pathways were ranked by −log10 of the adjusted P values. Pathways with significant enrichment (adjusted P ≤ 0.05) were further examined for relevance to glioma biology and brain/CNS-associated processes.

### Statistical Testing Analysis

Statistical comparisons were performed using appropriate tests based on data distribution and sample characteristics. One-way ANOVA was used to assess differences among sequencing pools. Given the non-normal distribution of most continuous variables, comparisons between glioma grades (HGG and LGG) were conducted using the Mann-Whitney U test. All statistical tests were defined as statistically significant if calculated p-values were less than or equal to 0.05.

GFs expressed in *Drosophila* were characterized quantitatively by their phenotypic effects on the VNC in terms of area (μm^2^), length (μm), area/length ratio, and glial cell migration distance (μm). For each phenotype, measurements from fusion-expressing fly lines were compared to wild-type (WT) controls using one-way ANOVA. Dunnett’s multiple comparison test was used to identify which fusions differed significantly from WT. Statistical analyses were performed using GraphPad Prism, data are presented as mean ± SEM, and statistical significance was defined as P ≤0.05.

## DISCUSSION

Long-read whole-transcriptome sequencing enabled comprehensive detection and characterization of GFs in glioma samples that were previously tested negative using the clinical CHOP FUSIP. In this fusion-negative cohort, the unbiased long-read approach identified numerous additional GFs, including those involving genes in the CHOP FUSIP and COSMIC Cancer Gene Census, that were not initially reported by the clinical test. This increased detection reflects the broader search of an untargeted whole-transcriptome assay and the ability of long-read sequencing to span entire fusion transcripts and resolve complex breakpoints.^62,63^ While short-read targeted transcriptome sequencing is also capable of identifying novel GFs, it typically infers the fusion structure from fragmented reads spanning breakpoint regions. In contrast, long-read sequencing directly captures full-length fusion transcripts, providing greater confidence in fusion structure, orientation, and isoform composition. However, current targeted short-read fusion panels and whole-transcriptome assays remain highly accurate and scalable for detecting known, recurrent fusions in routine clinical practice.^22^ Because this study compares a targeted short-read panel to an unbiased long-read whole-transcriptome approach (and does not include unbiased short-read RNA-seq), the increased detection we observe reflects both the broader search space and the capacity of long-reads to sequence full-length fusion transcripts.

Focusing on GFs that contained breakpoints within CDS/near exon boundaries, an arrangement that increases the likelihood of altering protein structure or regulation, these events showed strong enrichment for oncogenes, tumor suppressors, glioma-associated genes, and recurrent CNS-related fusion partners. Nearly all fusions involved at least one gene listed in the COSMIC Cancer Census, a small subset involved genes in the targeted panel, and over half included genes previously reported as recurrent fusion partner genes in CNS-related tumors, highlighting the involvement of genes with established roles in glioma-related oncogenesis and brain tumors. We also found that many fusions overlapped with genes with high dependency scores in glioma-related cell lines, suggesting that some of these partners are important for tumor survival.^50,51^ These findings illustrate how expanding beyond a fixed targeted gene panel can uncover likely clinically relevant fusions in tumors previously classified as fusion-negative. Long-read sequencing further allows precise characterization of breakpoint configurations of fusions, particularly when combined with orthogonal experimental validation at base-pair resolution.

In parallel, at the transcriptomic-level, long-read sequencing revealed widespread isoform-level dysregulation in glioma, including novel promoter usage, alternative splice junctions, and altered transcription termination and initiation sites, resulting in multiple NIC and NNIC novel isoforms across the glioma samples. Genes with high numbers of novel isoforms frequently displayed alternative first exons or non-canonical transcription start sites, consistent with enhancer dysregulation and promoter rewiring observed in glioma and other CNS tumors.^64^ Several genes with multiple novel isoforms, such as *BCAN*, *NDRG2*, and *TRPT1*, have been implicated in glioma biology or prognosis.^17,36,65^ Notably, for a subset of genes involved in GFs, we observed non-canonical transcript start or stop sites proximal to the reported fusion breakpoints, highlighting an association between local transcript architecture and fusion breakpoint localization. Genes with recurrent novel isoforms were enriched in pathways related to synaptic signaling, glial biology, and cell-cycle regulation, suggesting that isoform alterations may contribute to both oncogenic signaling and neural tissue development.

We further evaluated the oncogenic potential of selected GFs using functional assays in *Drosophila*. For these assays, patient-specific fusion constructs were generated based on breakpoint-defined open reading frames (ORFs) mapped to reference transcripts. Overexpression of selected GF ORFs in *Drosophila* glial cells produced a spectrum of phenotypes, including altered VNC development leading to its elongation/contraction and/or enlargement/reduction, and aberrant glial cell migration. These phenotypes are consistent with tumor-like overgrowth and infiltration described in established *Drosophila* glial tumor models^29^ and conceptually parallel the diffuse/infiltrative behavior observed in mammalian glioma models driven by kinase fusions such as *NTRK* and *FGFR* gene rearrangements.^66^ Not all fusion constructs produced measurable phenotypes, underscoring the likelihood that additional factors regulate fusion-driven oncogenicity. Nevertheless, the fusions that did elicit VNC phenotypes provide *in vivo* evidence that a subset of breakpoint-defined GFs that were identified in FUSIP-negative gliomas can affect glial development. These functional assays therefore support the oncogenic potential of selected GFs and highlight that long-read fusion discovery coupled with downstream *in vivo* screening can be leveraged toward large-scale precision oncology. This combined strategy that spanned discovery, transcript-level characterization, and functional validation, establishes a framework for refining fusion prioritization, identifying potentially targetable pathways, and guiding future incorporation of newly emerging fusions into clinical testing algorithms.

Although this study expands our understanding of glioma biology through long-read transcriptomics, there are a couple of important limitations that should be noted. The modest cohort size (49 glioma and 22 control samples) limits statistical power for detecting rare and grade-specific differences. Sequencing depth also varied across samples, potentially reducing sensitivity for low-abundance isoforms. This variability, together with modest sample size, may explain the lower number of significant isoforms in the HGG vs. LGG comparison. While a subset of novel GFs was functionally validated, most candidates were not experimentally tested. This is typical for large-scale structural variant and fusion discovery studies, where only a selected portion of predicted events can feasibly be validated, so we deliberately prioritized fusions with CDS/near exon breakpoints and strong cancer gene annotations for follow-up functional assays. As matched whole-transcriptome short-read RNA-seq data were not available, the increased fusion detection cannot be directly attributed solely to long-read sequencing. Instead, our findings demonstrate the combined effect of an unbiased transcriptome-wide assay and long-read resolution. Future benchmarking studies comparing matched long- and short-read whole-transcriptome data would be required to distinguish specific performance advantages attributable to long reads alone. Lastly, because matched genomic DNA was unavailable, somatic versus germline fusion events could not be confirmed. Although the absence of these GFs in the control brain tissues support their likely somatic origin, matched genomic analyses would have provided a more definitive patient-specific validation. Despite these constraints, the combined use of multiple GF and transcript isoform detection tools, complementary statistical approaches, and experimental functional validation provided high-confidence insights into glioma transcriptomic complexity. This integrated long-read sequencing and analysis framework enables systematic discovery and prioritization of candidate GFs beyond targeted panels, providing a foundation for multi-omic studies, functional validation, and future diagnostic or therapeutic development.

## CONCLUSION

This study demonstrates the power of ONT long-read whole-transcriptome sequencing to uncover potentially novel and clinically relevant GFs that were overlooked by the targeted short-read fusion panel used in this cohort. By integrating long-read sequencing with GF detection, transcript isoform analysis, and downstream functional validation, we identified and experimentally validated potentially novel GFs with pathogenic potential. Transcript-level alterations with non-canonical stop and new start sites in fusion partner genes provided additional context around the breakpoint regions and were consistent with fusion-associated alterations. Differential isoform expression and pathway analyses revealed similar changes across glioma grades, suggesting common pathways underlying disease progression. Functional assays in *Drosophila* revealed tumor-like phenotypes in glial cells that are consistent with altered growth and tumor progression. Together, these findings demonstrate that long-read whole-transcriptome sequencing expands GF detection and transcript isoform profiling and, when coupled with *in vivo* functional validation in *Drosophila*, provides new biological insights into glioma that supports advancing precision diagnostics for fusion-negative gliomas.

## DATA AVAILABILITY

ONT long-read sequencing data for both glioma sample cohorts have been deposited at the NCBI Sequence Read Archive (SRA) under the BioProject number PRJNA1267462, which are publicly available as of the date of publication.

## AUTHOR CONTRIBUTIONS

Conceptualization, K.W., Y.S., M.L., and K.R.; Methodology, K.W., Y.S., M.L., K.R., M.U.A., and H.M.D.; Formal Analysis: K.R., E.N.Y.C., E.G., M.U.A., and H.M.D.; Investigation: K.R., E.G., E.N.Y.C., H.M.D., F.X., and J.C.; Resources, K.W., Y.S., and M.L.; Data Curation, K.R., E.G., M.U.A., J.C., E.N.Y.C., and H.M.D.; Writing – Original Draft, K.R., E.N.Y.C.; Writing – Review & Editing, K.R., E.N.Y.C., H.M.D., M.U.A., E.G., K.W., Y.S., and M.L.; Visualization, K.R. and E.N.Y.C.; Supervision, K.W., Y.S., and M.L.; Funding Acquisition, K.W., Y.S., and M.L.; All authors reviewed the manuscript.

## ACKNOWLEDGEMENTS

The authors would like to thank our collaborators at the Division of Genomic Diagnostics (DGD) at The Children’s Hospital of Philadelphia (CHOP) and the University of Pennsylvania. We thank technical consultation from the IDDRC Biostatistics and Data Science core (HD105354). This study is supported in part by NIH grant GM132713 and HG013359, a NIH/NIHGRI T32 training grant on Computational Genomics (HG000046), and a CHOP Omics Initiative grant.

## Notes

### Competing Interest Statement

The authors have declared no competing interest.

