## Supplementary Information for "Long-Read Transcriptome Sequencing and Functional Validation Reveals Novel and Oncogenic Gene Fusions in Fusion Panel-Negative Gliomas"

*SUPPLEMENTAL INFORMATION*

**Supplemental Figure S1. Computational and experimental validation of additional candidate GFs. (A)** IGV plot (top) with a subset of the input reads (gray), highlighting split alignments within the chromosomal regions of interest. Gene names and reported breakpoints (indicated by dotted lines) are annotated for clarity. Genome Ribbon single-read view (bottom) with labeled gene regions by name and the input read orientation is denoted by the arrow. GFs shown in **(A)**: **(A1)** *ABL1::WDR83OS* (Sample 38), **(A2)** *BCL11A::VIRMA* (Sample 48), **(A3)** *CTNND2::EIF5A* (Sample 34), **(A4)** *MDM2::NUP107* (Sample 64), **(A5)** *PRICKLE1::FHIT* (Sample 40), **(A6)** *DDX6::DCTN2* (Sample 64). **(B)** Gel electrophoresis validation. The K562 cell line was used as a negative control, and validation was performed using RNA extracted from patient samples as noted. GFs shown in **(B)**: **(B1)** *ABL1*::*WDR83OS*, *BCL11A*::*VIRMA*, *MDM2::NUP107, PRICKLE1*::*FHIT*, *DDX6*:*DCTN2*. **(B2)** *CTNND2*::*EIF5A*, *DDX6*:*DCTN2* (*re-designed primers), *MDM2::NUP107* (*re-designed primers)*,* *PRICKLE1*::*FHIT* (*re-designed primers). **(C)** Sanger sequencing traces of positively validated GFs. GFs shown in **(C)**: **(C1)** *ABL1*::*WDR83OS* and **(C2)** *MDM2:: NUP107*.

**A1.**

**Sample 38**
***ABL1*::*WDR83OS* Fusion**

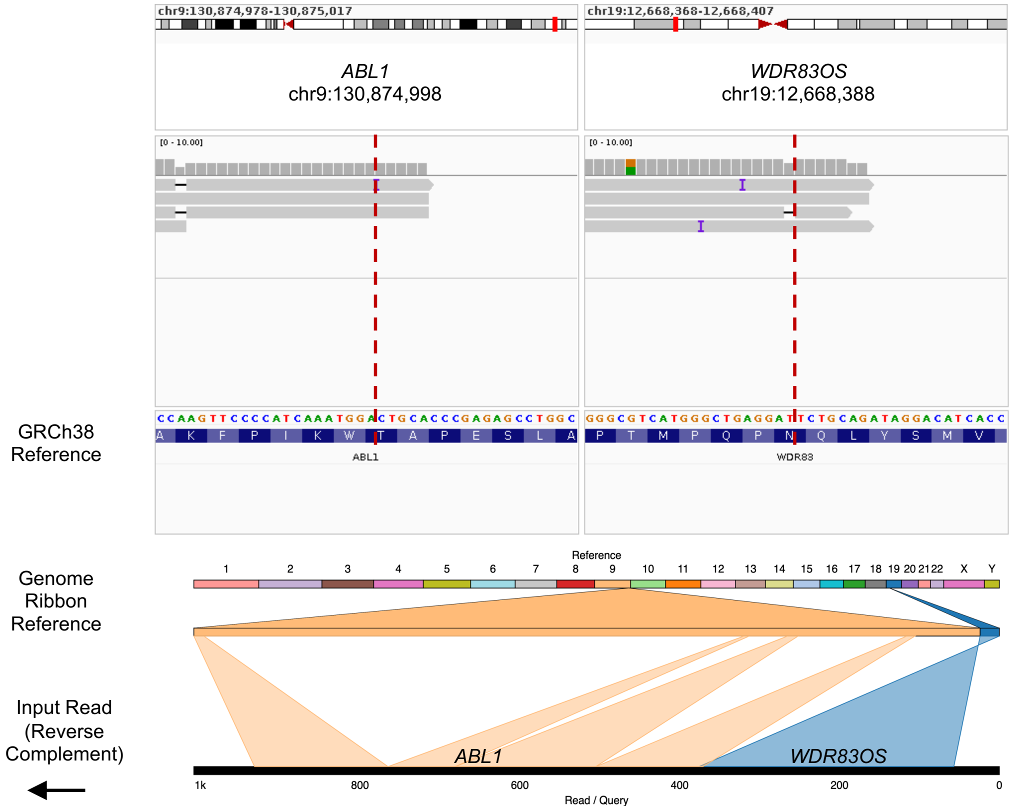

**A2.**

**Sample 48**
***BCL11A*::*VIRMA* Fusion**

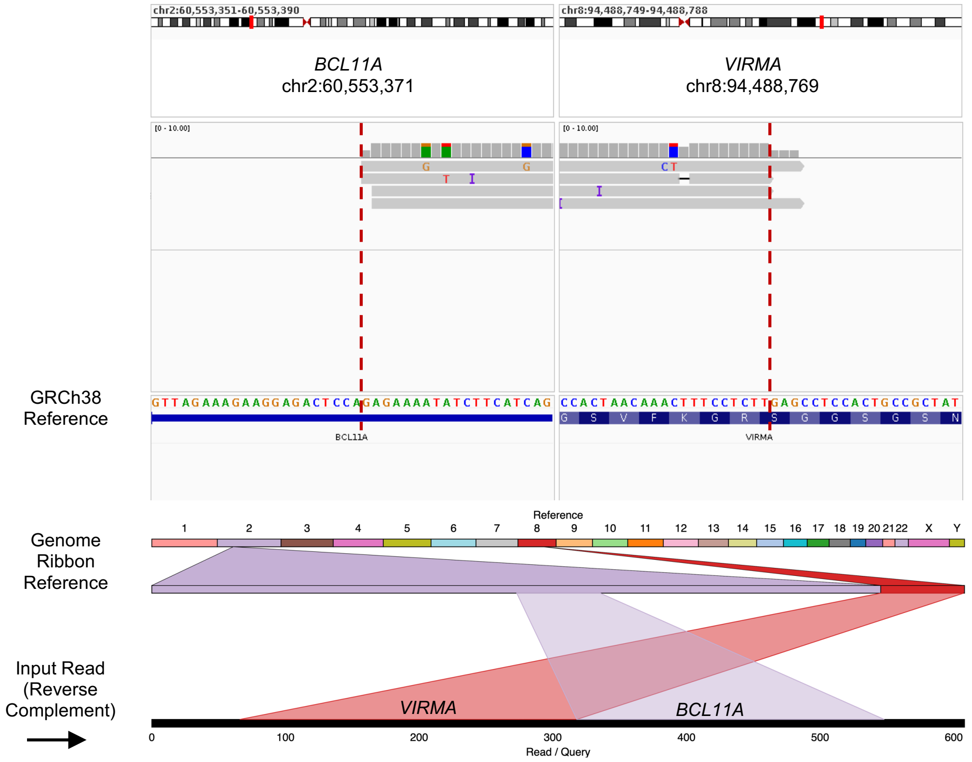


**A3.**

**Sample 34**
***CTNND2*::*EIF5A* Fusion**

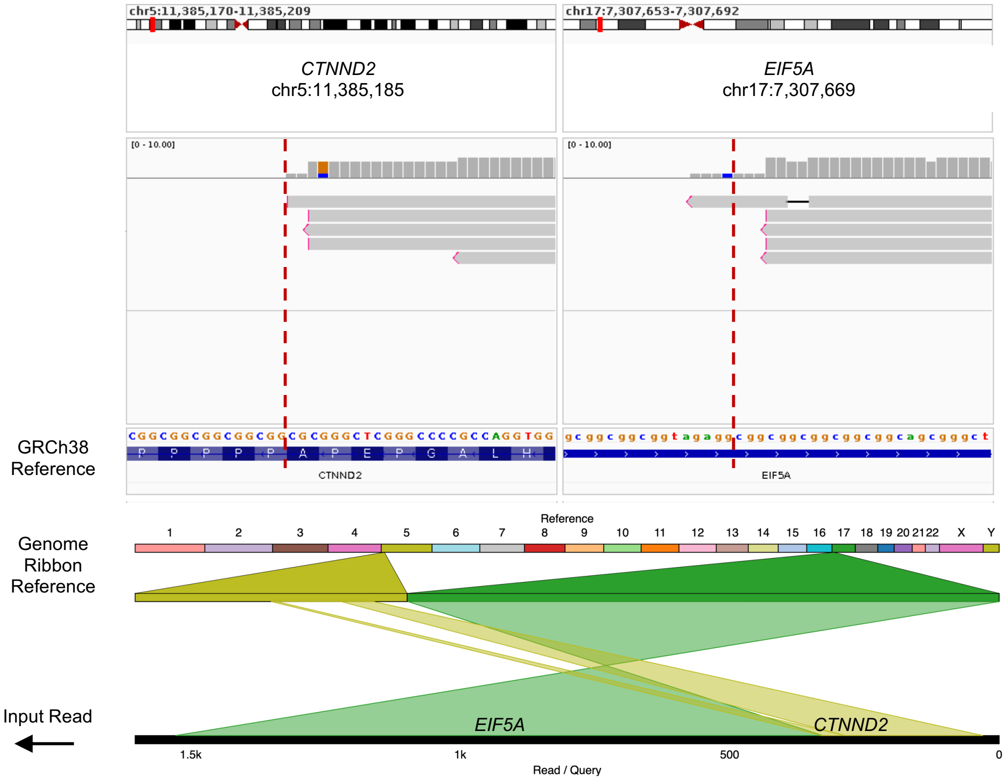

**A4.**

**Sample 64**
***MDM2::NUP107* Fusion**

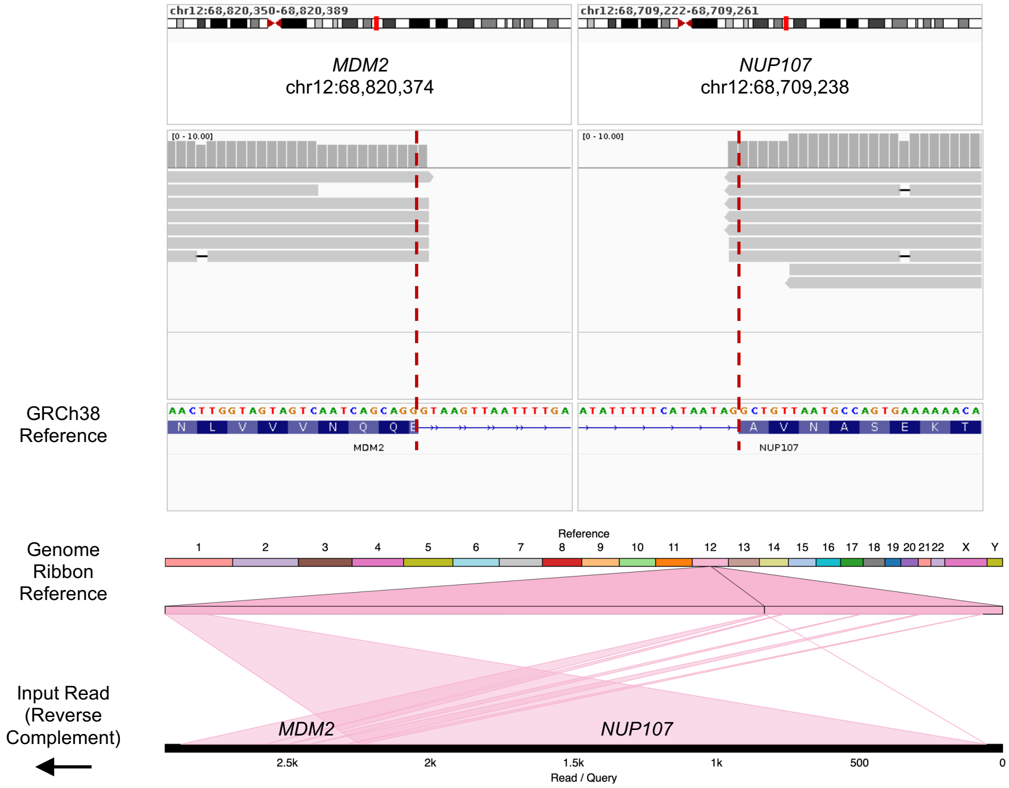


**A5.**

**Sample 40**
***PRICKLE1::FHIT* Fusion**

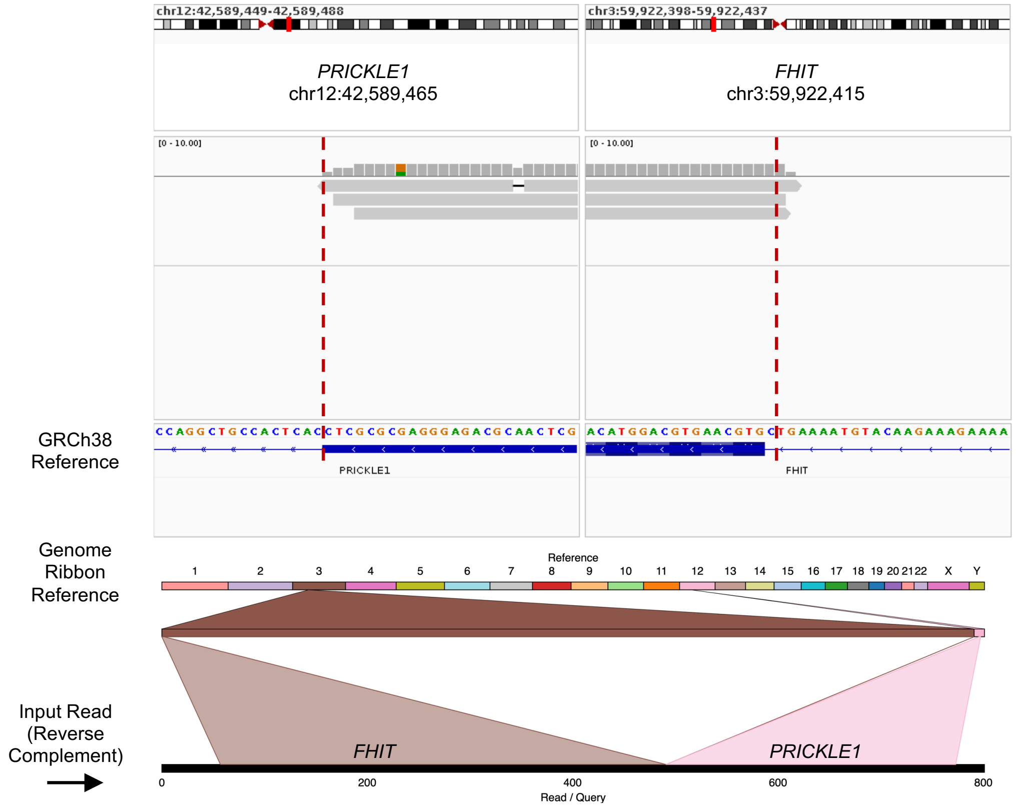

A6.

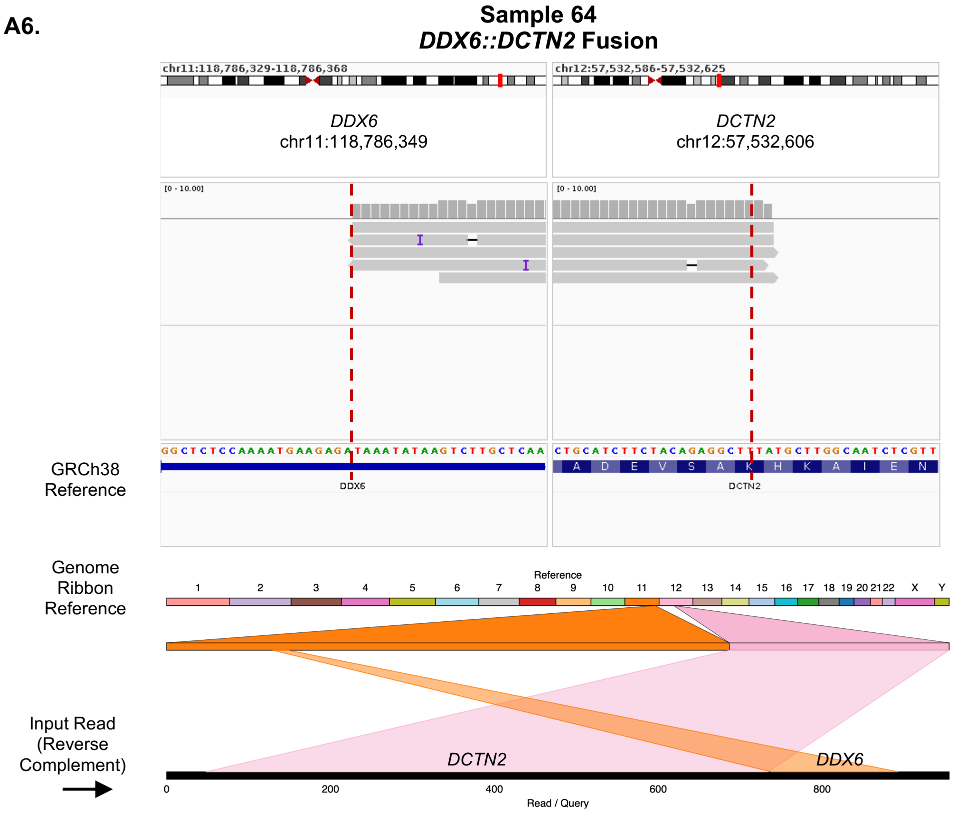
Sample 64
DDX6::DCTN2 Fusion
chr11:118,786,329-118,786,368
chr12:57,532,586-57,532,625
DDX6
chr11:118,786,349
DCTN2
chr12:57,532,606
[0 - 10.00]
[0 - 10.00]
GRCh38 Reference
GGCTCTCCAAAATGAAGAGATAAATATAAGTCTTGCTCAA
CTGCATCTTCTACAGAGGCTTTATGCTTGGCAATCTCGTT
A D E V S A K H K A I E N
DDX6
DCTN2
Genome Ribbon Reference
Reference
1 2 3 4 5 6 7 8 9 10 11 12 13 14 15 16 17 18 19 20 21 22 X Y
Input Read (Reverse Complement)
DCTN2
DDX6
0 200 400 600 800
Read / Query

B1.

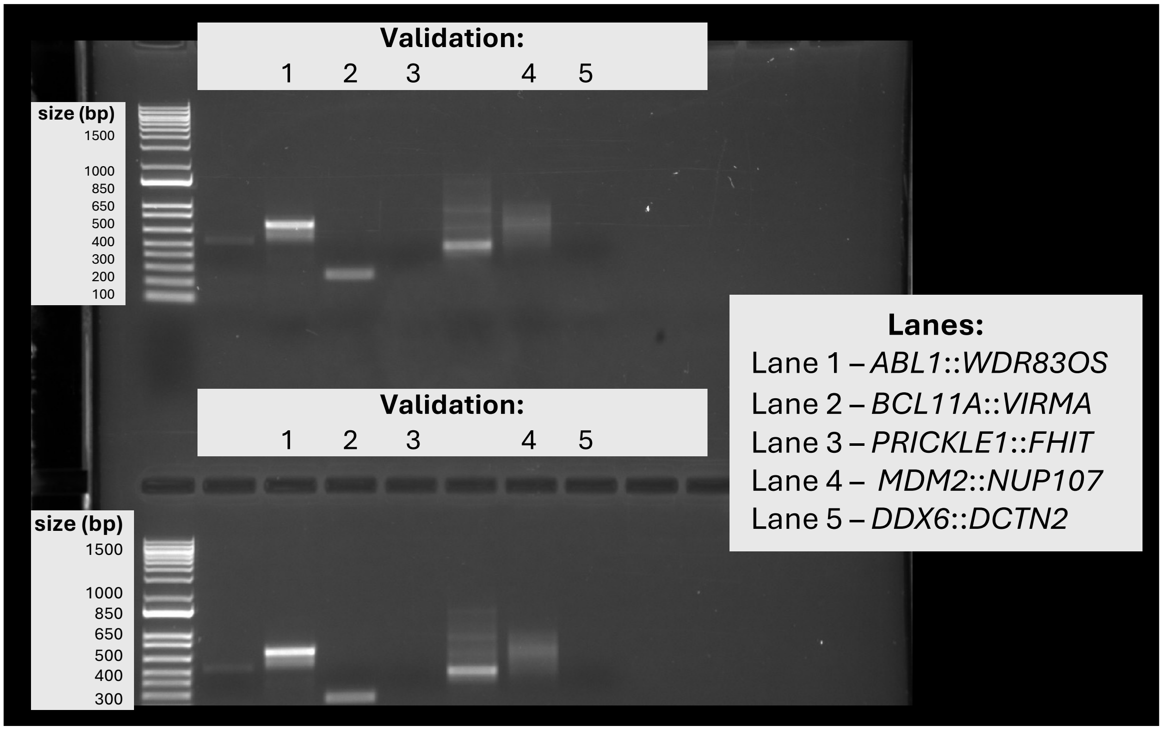
Validation:
1 2 3 4 5
size (bp)
1500
1000
850
650
500
400
300
200
100
Validation:
1 2 3 4 5
size (bp)
1500
1000
850
650
500
400
300
Lanes:
Lane 1 – ABL1::WDR83OS
Lane 2 – BCL11A::VIRMA
Lane 3 – PRICKLE1::FHIT
Lane 4 – MDM2::NUP107
Lane 5 – DDX6::DCTN2

**B2.**

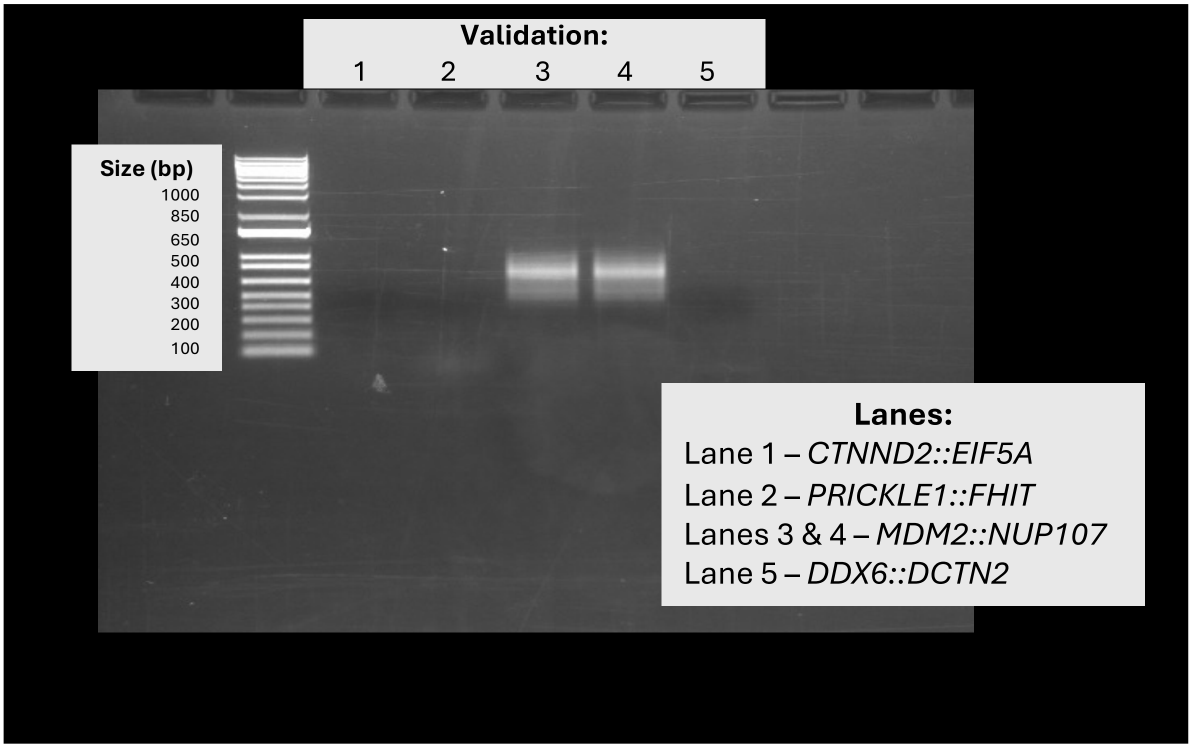


**C1.** *ABL1::WDR83OS*

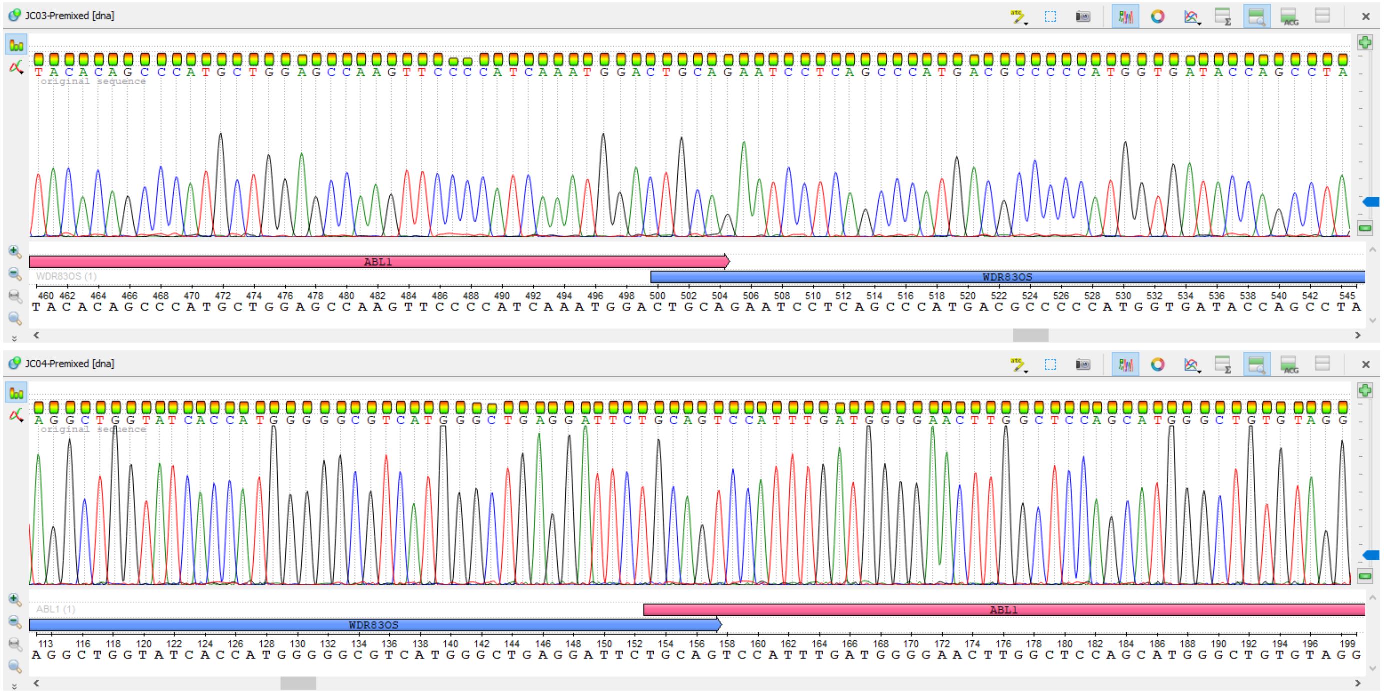

**C2.** *MDM2::NUP107*

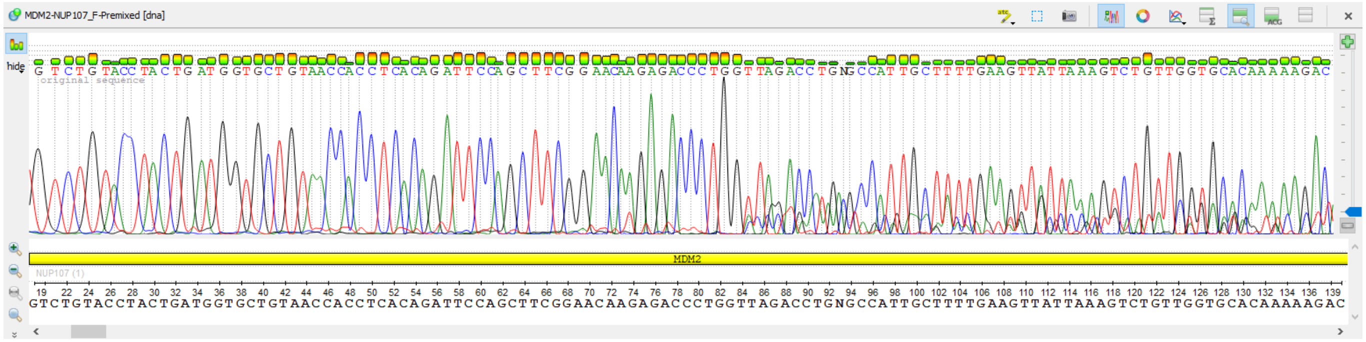


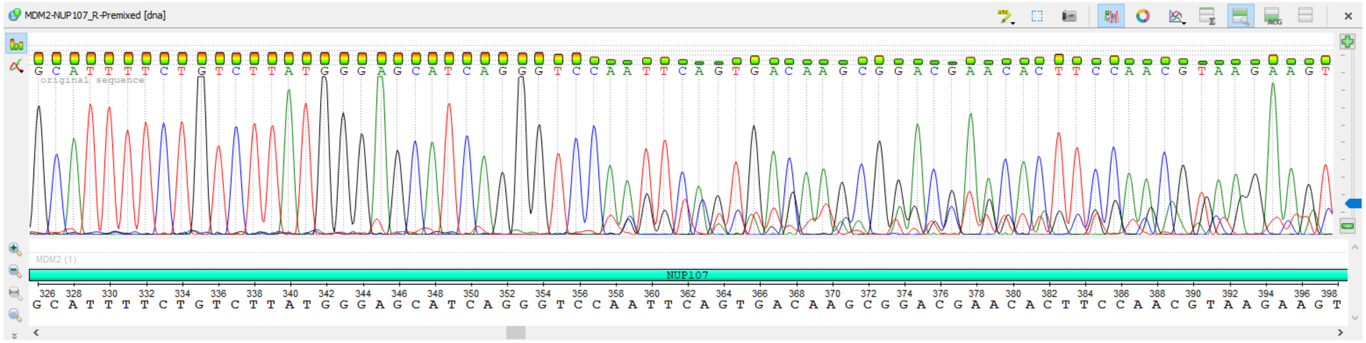

**Supplemental Figure S2. Differential transcript isoform expression across glioma comparisons for IsoQuant and FLAIR derived transcripts.** Concordance scatter plots between edgeR and DESeq2 are shown for all four glioma comparisons (Glioma vs. Controls, HGG vs. Controls, LGG vs. Controls, and HGG vs. LGG) for isoforms identified by IsoQuant **(A)** and FLAIR **(B)**. Each point represents an individual transcript isoform, with edgeR log2 fold change on the y axis and DESeq2 log2 fold change on the x axis. Significant differentially expressed isoforms from both tools (DESeq2 (FDR in edgeR, adjusted p-value in DESeq2 ≤0.05, and the absolute value of the log2 FC in expression ≥1) are shown, where concordant isoforms (significant in both edgeR and DESeq2) are colored red, DESeq2 only are colored blue, edgeR only are colored green, and gray for not significant in either method. Concordant isoforms cluster along the dotted diagonal (y = x), illustrating strong agreement in effect sizes between tools across all comparisons.

**A. IsoQuant**

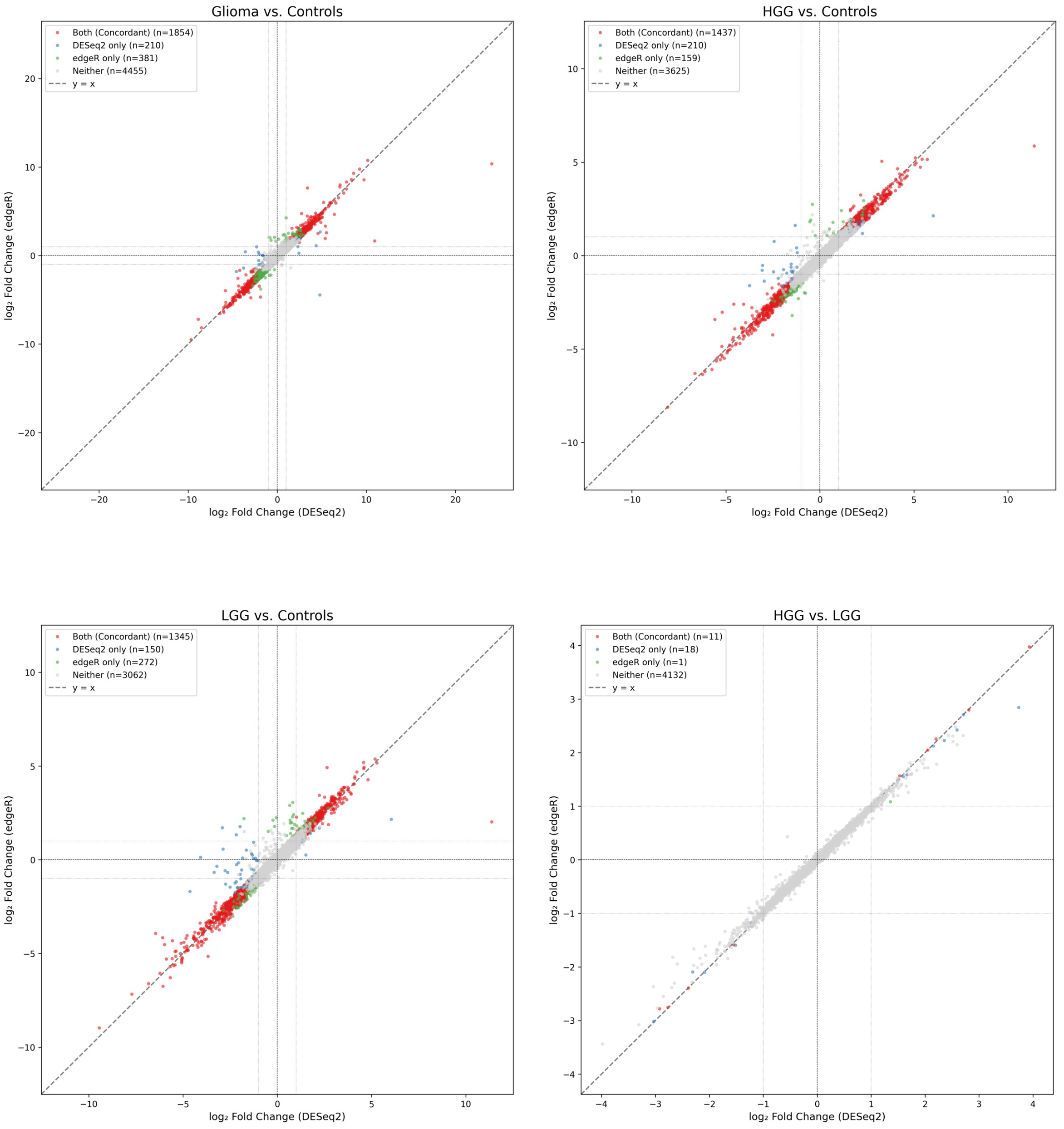

### B. FLAIR

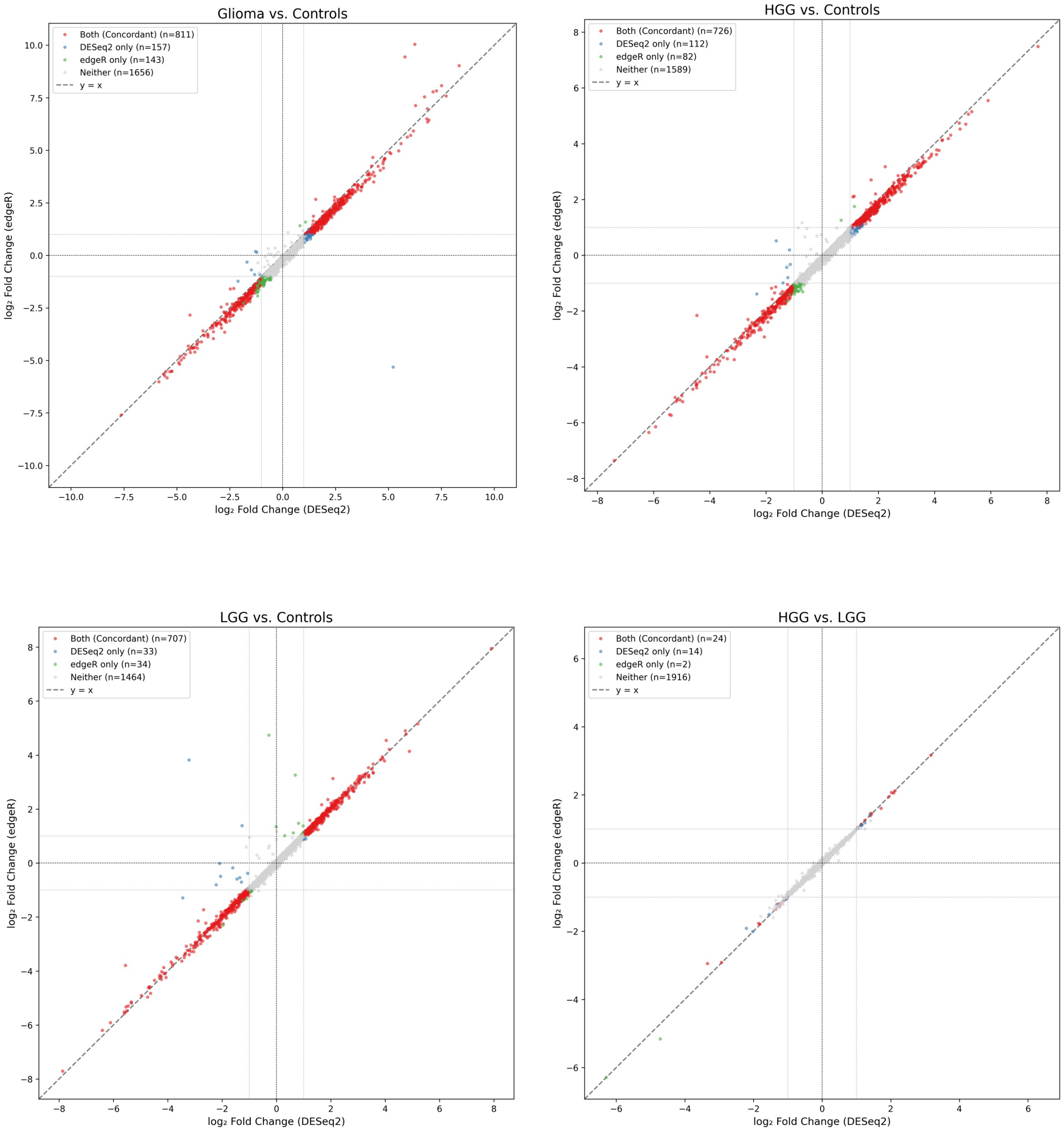

Glioma vs. Controls
Both (Concordant) (n=811)
DESeq2 only (n=157)
edgeR only (n=143)
Neither (n=1656)
y = x
log₂ Fold Change (DESeq2)
log₂ Fold Change (edgeR)
HGG vs. Controls
Both (Concordant) (n=726)
DESeq2 only (n=112)
edgeR only (n=82)
Neither (n=1589)
y = x
log₂ Fold Change (DESeq2)
log₂ Fold Change (edgeR)
LGG vs. Controls
Both (Concordant) (n=707)
DESeq2 only (n=33)
edgeR only (n=34)
Neither (n=1464)
y = x
log₂ Fold Change (DESeq2)
log₂ Fold Change (edgeR)
HGG vs. LGG
Both (Concordant) (n=24)
DESeq2 only (n=14)
edgeR only (n=2)
Neither (n=1916)
y = x
log₂ Fold Change (DESeq2)
log₂ Fold Change (edgeR)

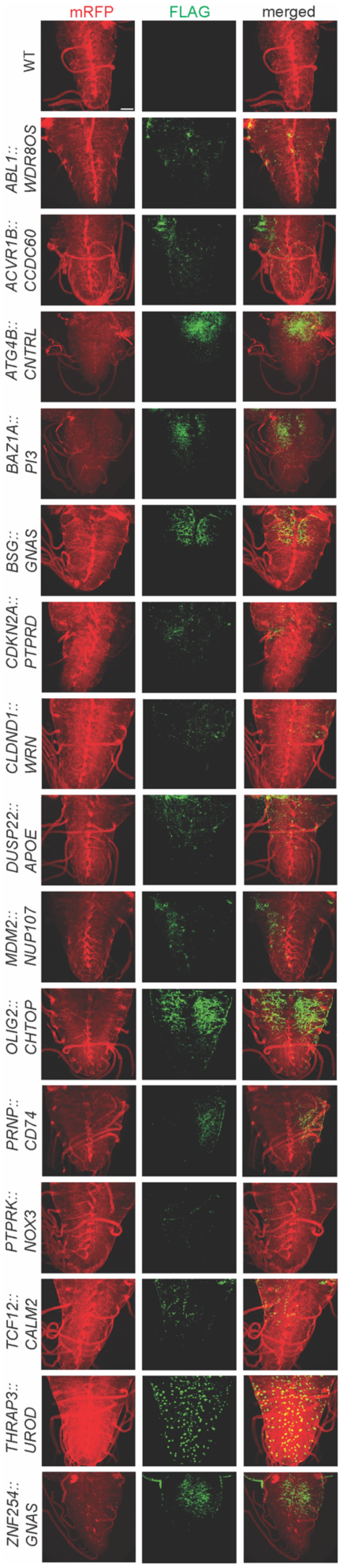

mRFP
FLAG
merged
WT
ABL1:: WDR8OS
ACVR1B:: CCDC60
ATG4B:: CNTRL
BAZ1A:: PI3
BSG:: GNAS
CDKN2A:: PTPRD
CLDND1:: WRN
DUSP22:: APOE
MDM2:: NUP107
OLIG2:: CHTOP
PRNP:: CD74
PTPRK:: NOX3
TCF12:: CALM2
THRAP3:: UROD
ZNF254:: GNAS

**Figure S3. FLAG immunostaining confirms expression of breakpoint-defined GFs in *Drosophila melanogaster* transgenic lines.** Confocal images of dissected third instar larval VNCs with *repo-GAL4* driving *mRFP* expression and GFs in glia. GF expression was detected by immunostaining of the FLAG epitope (green channel). All GFs show detectable FLAG signal, with localization observed in glial processes and/or nuclei. Scale bar = 50 μm.
